# Brainwide input-output architecture of paraventricular oxytocin and vasopressin neurons

**DOI:** 10.1101/2022.01.17.476652

**Authors:** S.N. Freda, M.F. Priest, D. Badong, L. Xiao, Y. Liu, Y. Kozorovitskiy

## Abstract

Oxytocin and vasopressin are pleiotropic neuropeptides with well-established roles in the regulation of social behavior and homeostatic functions. Their structural similarity and conserved functions in vertebrate social behavior suggest that neurohypophyseal peptides may represent a single integrative neuromodulatory system, yet both peptides subserve sexually dimorphic functions at the behavioral level. The extent to which central oxytocin and vasopressin systems share similar circuit architecture has not been previously studied. Sex differences in the central circuitry of the oxytocin and vasopressin systems may underlie sex-variant behaviors, but it is currently unknown whether the synaptic inputs or outputs of each neuropeptidergic system vary across males and females. To close this gap, we generated quantitative anterograde and retrograde maps of the paraventricular oxytocin and vasopressin systems in mice. We observed that both oxytocinergic and vasopressinergic neurons share highly similar synaptic inputs that are sex-conserved. Projection patterns differed across systems and showed sex differences, more pronounced in the vasopressin neurons. Together our data represent the first comparative study of oxytocin and vasopressin input-output architecture highlighting how these neurohypopheseal peptides can play complementary and overlapping roles that are sex-dependent.

## Introduction

Oxytocin and vasopressin are highly conserved neuropeptides that underlie complex social behavior and physiological mechanisms^1–3^. These pleiotropic peptides are composed of nine amino acids and their molecular structures differ at only two amino acid positions. It is hypothesized that oxytocin and vasopressin originated from a local gene-duplication event and have evolved into an integrative system necessary for reproduction, defensive behavior, and social attachment^4^. The behavioral effects of oxytocin and vasopressin are largely sex dependent. In female rodents, oxytocin is critical for maternal behavior, lactation, and sociosexual approach^5–8^, while in males, oxytocin release underlies consummatory sexual behaviors, modulates aggression, and reduces anxiety-related behaviors^9–12^. Vasopressin similarly exhibits sexual dichotomy in its behavioral mechanisms, with different effects on stress adaptation, defensive behaviors, and pair bond formation in male and females^13–15^. Yet, how these behavioral differences are produced is still not fully resolved.

Separate populations of neurons in the hypothalamus synthesize oxytocin and vasopressin. Oxytocin and vasopressin-producing neurons primarily reside in the paraventricular nucleus of the hypothalamus (PVN) and supraoptic nucleus (SON), with an additional small population of vasopressin-synthesizing neurons in the suprachiasmatic nucleus. Together, these neurosecretory cells send projections throughout the brain to modulate circuits for social behavior and facilitate peripheral release through contacts to the posterior pituitary gland^3,16^. Sensory experience alters oxytocin and vasopressin neuronal activity—pup distress calls, social touch, and observational learning of pup retrieval have all been shown to activate and change PVN oxytocin neuron firing^6,17–19^. While less is known about social experience-dependent changes in vasopressin neurons, stressors and fluctuations in blood osmolarity induce activity in vasopressin neurons and elicit vasopressin peripheral release. To fully understand how oxytocin and vasopressin can subserve disparate behaviors and function cooperatively to promote sociality, we must delineate the circuit architecture—the axonal projections and presynaptic inputs—of each system.

Sex-specific behaviors of the oxytocin and vasopressin systems are driven by differences in either the central or peripheral nervous system. Oxytocin and vasopressin receptor distributions differ in male and female rodent brains^20,21^, yet whether differences occur in the inputs and outputs of these neuronal populations is currently not reported. To understand if sex-variant behaviors are due to differences in the central nervous system, we performed brain-wide quantitative anterograde and retrograde tracing of paraventricular oxytocin and vasopressin neurons in mice. We used a modified version of the open-source software, WholeBrain^22^, to register 2D coronal slices to the Allen Institute mouse reference atlas^23^. We find that the circuit architecture of the oxytocin system is largely sex-invariant, while vasopressin projection patterns appear sex-dependent. Together, our data compiles the first comparative and quantitative input-output map of paraventricular oxytocin and vasopressin neurons, highlighting how the central architecture of these pleiotropic peptides may subserve complementary or opponent socioregulatory behaviors.

## Results

### Brainwide axonal projections of paraventricular nucleus oxytocin neurons

To generate a complete map of axonal projections from PVN oxytocin (Oxt) neurons and compare projection patterns across sexes, we performed quantitative brainwide mapping of axonal projections of Oxt PVN neurons in male and female mice. We injected a Cre-dependent eYFP (AAV9-EF1a-DIO-eYFP) into the PVN of Oxt ^*i-Cre*^ mice^24,25^ and acquired coronal slices through the entire mouse brain for quantitative 2D mapping (Figure 1 a-b, Figure 1-Figure Supplement 1a-b). This method offers high specificity in genetic targeting of Oxt neurons^26,27^. In addition to the PVN, a population of Oxt neurons also resides in the supraoptic nucleus (SON) of the hypothalamus. Oxt-synthesizing neurons in this nucleus send direct projections to the posterior pituitary gland to control peripheral release of Oxt. Since we were interested in centrally-projecting Oxt neurons, our analyses were focused on PVN neurons and excluded subjects exhibiting expression in the SON.

**Figure 1:**
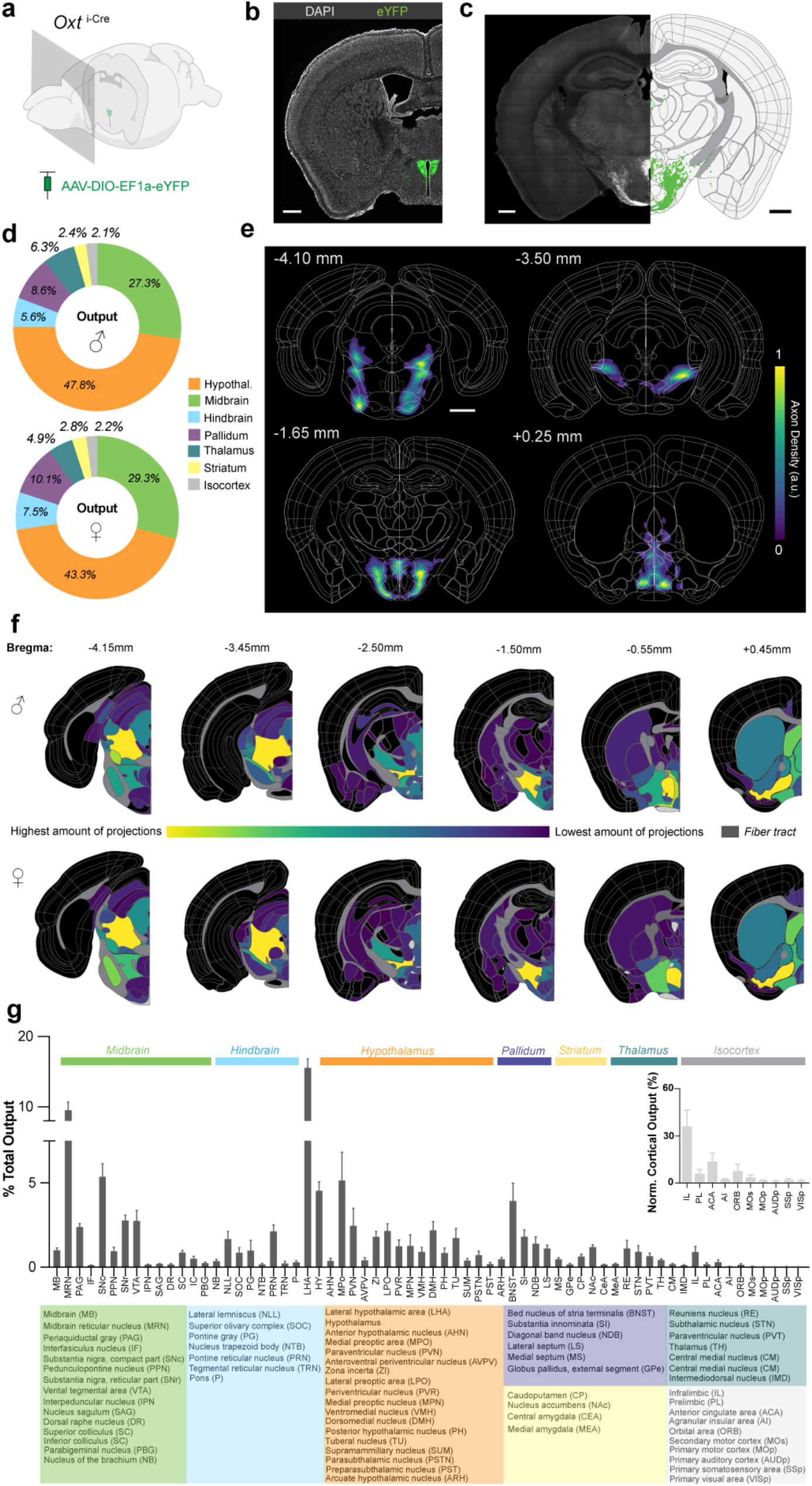
Brainwide mapping of Oxt+ projections from the PVN shows broad, sex-invariant targeting. a. Schematic of AAV9-DIO-eYFP injection into PVN of Oxt^i-Cre^ mice. Coronal sections were collected at an interval of 180 µm. b. Representative image of eYFP expression in Oxt+ PVN neurons. Scale bar, 0.5 mm. c. *Left* Example raw image of Oxt+ projections from the PVN to local hypothalamic regions. *Right* Registered coronal slice to the Allen Brain atlas, with segmented axonal projections of brain slice (*left*). d. Comparison of average total axonal output of Oxt+ neurons across male (n=3) and female (n=3) mice in parent regions of the Allen brain atlas: hypothalamus (hypothal.), midbrain, hindbrain, pallidum, striatum, and isocortex. The majority (>65%) of Oxt+ projections are in the midbrain and neighboring hypothalamic nuclei in both male and female mice. e. Projection density heatmaps of representative female mouse coronal sections. Regions with greatest density of projections are depicted in yellow; regions containing few projections, purple. Bregma coordinates are depicted above each registered section. Scale bar, 1 mm. f. Coronal heat maps of one male and one female brain. Each nucleus color corresponds to normalized projections. Nuclei with highest number of projections are in yellow; areas with little to no projections are in purple or black. Projection patterns are largely similar across male and female mice. g. Brain nuclei showing Oxt+ projections in male and female mice (n=6 total), quantified as % of total quantified projections in the brain. Each region depicted in graph contains >0.15 % of total projections, except for some cortical regions. *Inset* Total quantified projections normalized to isocortical nuclei. Medial prefrontal cortex and orbital cortex show most Oxt+ projections relative to other cortical regions. Brain region acronyms with corresponding parent regions are depicted below the bar graph. For a list of all brain areas quantified, see Table 1.

We registered brain slices to Allen Institute mouse reference atlas using the open-source R package, WholeBrain^22^ (Figure 1c, Figure 1-Figure Supplement 1c). 2D mapping of coronal sections to standardized brain atlases is simple, efficient, and cost-effective, relative to more complex approaches like 3D tomography or other volumetric processing which compress 3D data for analyses and visualization (Figure 1-Figure Supplement 1c). Axonal projections were identified via a Sobel edge detection algorithm and subsequently quantified as pixel counts in each registered brain region. For a subset of brains, we collected posterior cerebellar and hindbrain slices and found limited axonal projections in cerebellar cortex (<1% total output) and sparse projections (<0.1% total output) in the medulla that likely correspond to fibers extending to spinal cord^12,28^ (Figure 1-Figure Supplement 1d-e). Therefore, our data sets exclude these regions from the brain-wide mapping and focus solely on cerebral and brainstem nuclei.

We found PVN Oxt projections were widely dispersed throughout the mouse brain, extending as far as forebrain neocortical nuclei and posterior hindbrain regions, and were primarily localized to ventrally located nuclei (Figure 1d-f, Figure 1-Figure Supplement 2 and 3, Table 1). On average, 43% of Oxt axons were found in hypothalamic nuclei, with the largest proportion residing in the lateral hypothalamic area (LHA: 13.1±1.3% ♀, 17.9±1.1% ♂ total PVN Oxt+ output), zona incerta (ZI: 1.9 ±0.2% ♀, 1.7±0.7% ♂), and medial preoptic nucleus (6.8 ±3.0% ♀, 3.5±1.5% ♂, Figure 1g). Considering that the LHA is adjacent to the PVN, Oxt projections quantified in this region may be fibers of passage and not axon terminal fields. Midbrain nuclei comprise the next largest proportion of Oxt projections, contributing an average 30% of total Oxt axonal projections. This primarily includes the midbrain reticular nucleus (MRN: 9.7±2.3% ♀, 9.2 ±1.3% ♂), periaqueductal gray (PAG: 2.5±0.3% ♀, 2.3±0.3% ♂) and many basal ganglia nuclei critical for reward and salience processing, such as the substantia nigra pars compacta (SNc: 5.6±1.6% ♀, 5.2±0.7% ♂), ventral tegmental area (VTA: 2.5±0.9% ♀, 3.0±1.1% ♂), pedunculopontine nucleus (PPN: 1.1±0.5% ♀, 0.8±0.1% ♂), and substantia nigra reticulata (SNr: 3.0±0.6% ♀, 2.5±0.4% ♂).

**TABLE 1.**
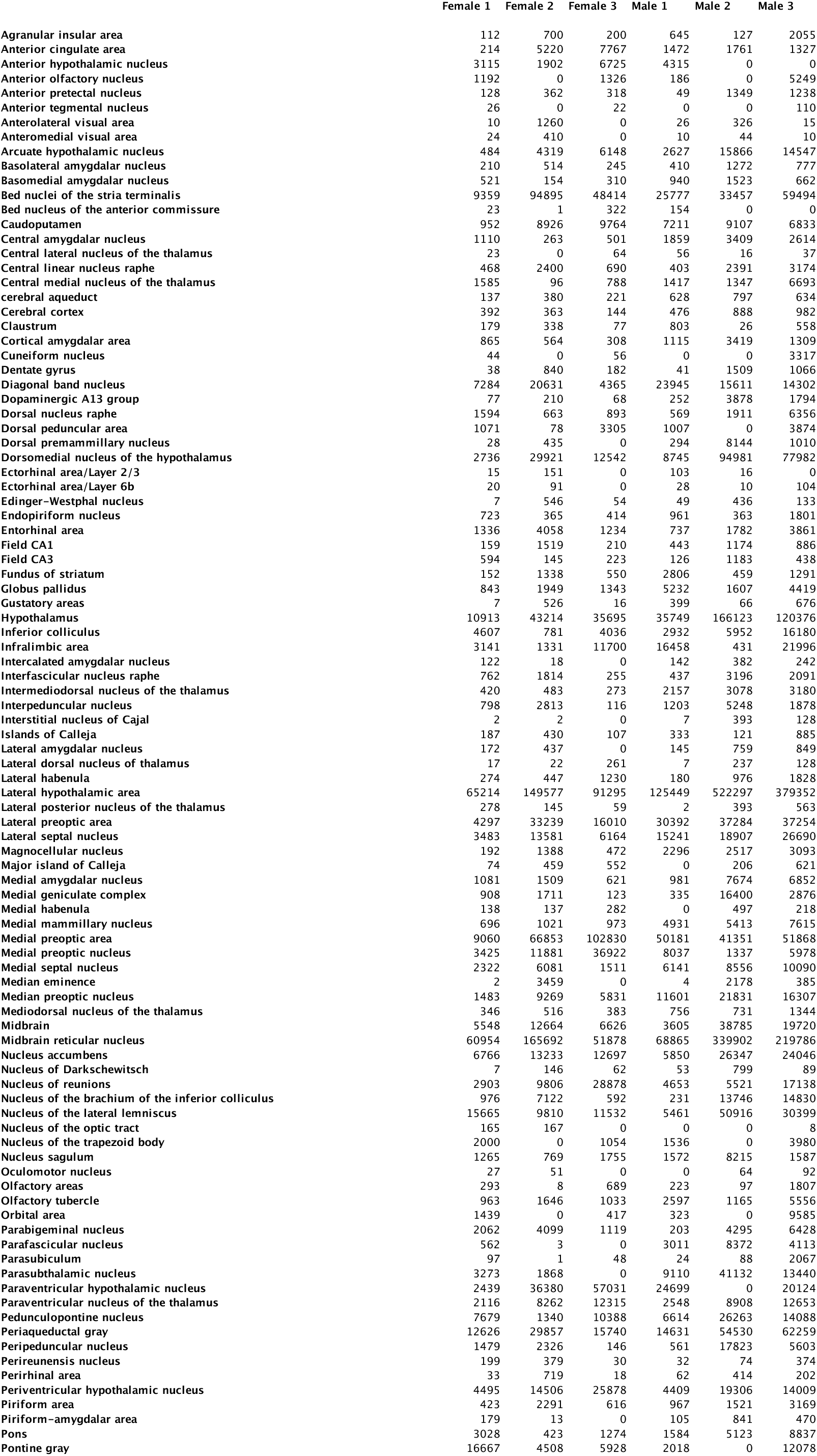
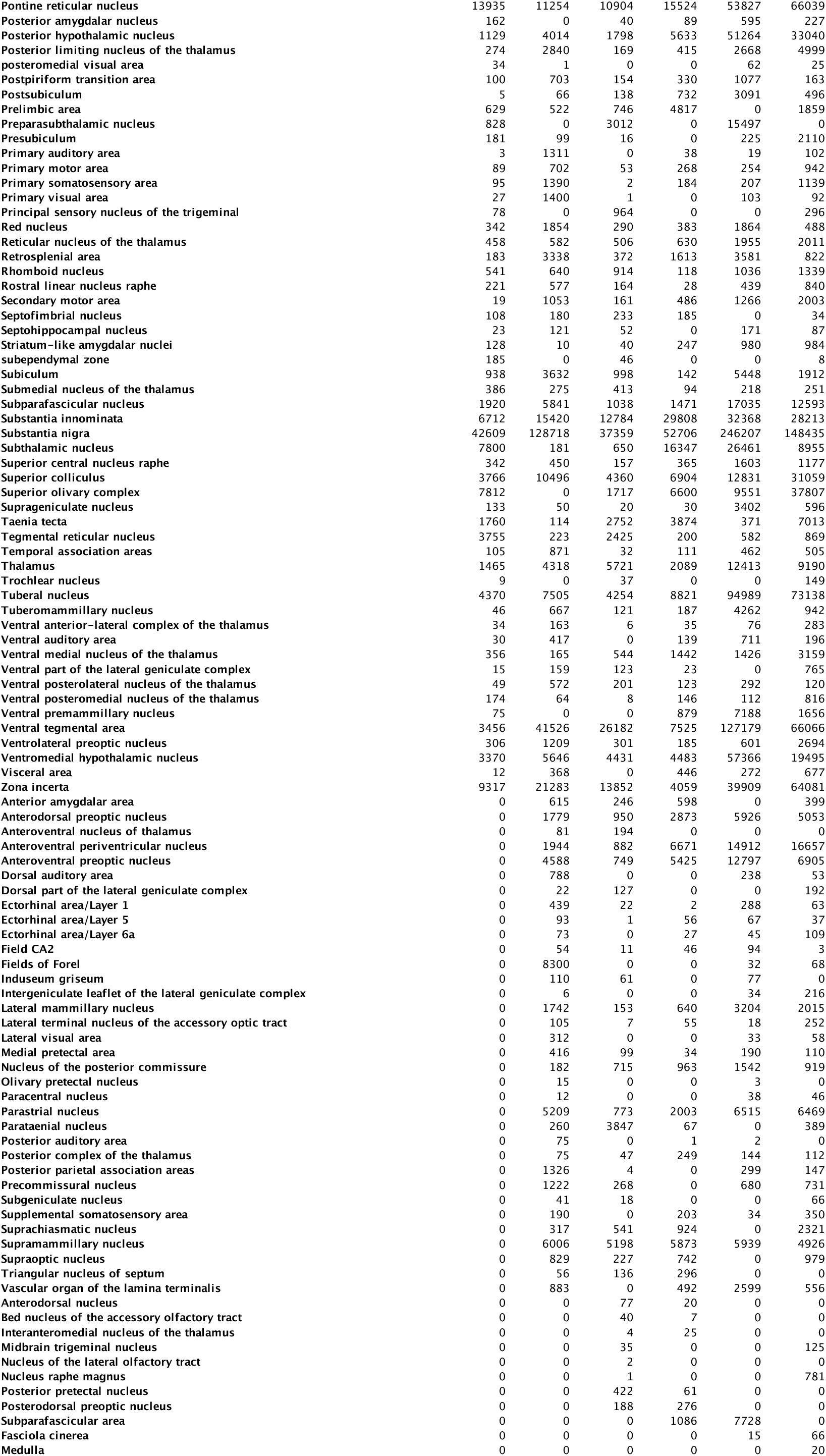
Total oxytocin projection pixel counts per sample (related to Figure 1)

More anterior striatal and pallidal regions exhibited less prominent Oxt axonal content by comparison. The bed nucleus of the stria terminalis (BNST: 5.4±1.7% ♀, 2.5±0.6% ♂), substantia innominata (SI: 1.5±0.1% ♀, 2.1±0.9% ♂), and nucleus accumbens (1.4±0.2% ♀, 1.0±0.1% ♂) contain some of the densest projections within these areas. Isocortical nuclei contain the smallest proportion of Oxt output, with an average of 2.2% of total axonal projections (Figure 1d). When axonal output is normalized to all cortical regions, the medial prefrontal cortex—particularly the infralimbic subregion—accounts for the majority of long-range Oxt cortical axons, even when compared to other cortical structures such as auditory cortex known for Oxt-mediated social behavior (Figure 1g)^29,30^. Finally, our data confirmed that PVN Oxt output is largely sex-invariant, with only two significant differences across males and females (Figure 1-Figure Supplement 4a-b, two-way ANOVA with Bonferroni adjustment for multiple comparisons). The two regions to show differences across males and females were the LHA (p<0.0001) and medial preoptic area (MPo, p=0.01). Given the proximity between PVN injection site and LHA, the difference in LHA axonal density may relate to fibers of passage and injection targeting, but we cannot exclude the possibility that it represents a biologically meaningful difference (Figure 1-Figure Supplement 2 and 3). The MPo is a well-known anatomically sexually dimorphic brain region in many species, and prior work describes sex differences in axonal distributions of other neuromodulators linked to social processing^31^. Here, the observed difference across males and females is small (p=0.0113, 6.8±3.0% ♀, 3.5±1.5% ♂). In sum, differences in Oxt-regulated behavioral output are likely not due to variance in gross anatomical output architecture of the Oxt system in males and females. Our data also highlight new regions where Oxt release is currently understudied, such as the periaqueductal gray, midbrain reticular nucleus, and pedunculopontine nucleus.

### Retrograde input mapping of paraventricular nucleus oxytocin neurons

Control of Oxt release by inputs to centrally-projecting PVN neurons presents another potential mechanism for sexually dimorphic effects of Oxt on behavior. Prior anatomical and electrophysiological studies suggest that Oxt release from the PVN is primarily regulated by input from local GABAergic neurons from neighboring hypothalamic nuclei, including the lateral hypothalamic nucleus and medial preoptic area^32–35^. While multiple prior reports support the existence of inputs to Oxt neurons from neighboring hypothalamic regions, more recent studies have raised the possibility of longer-range projections from distal nuclei such as the superior colliculus and septal nuclei^19,36^. To date, a brainwide input map of Oxt neurons for both sexes has not been published. In order to map synaptic inputs to Oxt+ neurons and determine whether they differ across male and female mice, we performed genetically targeted retrograde mapping using pseudotyped rabies virus tracing (Figure 2a, Figure 2-Figure Supplement 1a). Cre-dependent helper virus expressing GFP was injected into the PVN, which was subsequently transduced with rabies virus strain RVdG fused with mCherry (Figure 2b, Figure 2-Figure Supplement 1a-b). Starter cell locations and counts did not vary across sex (Fig 2c: 44.8±6.8 cells ♀, 57.5±18.1 cells ♂, p=0.91 Mann-Whitney t-test). Starter cells were excluded from quantification of total inputs to PVN Oxt neurons.

**Figure 2:**
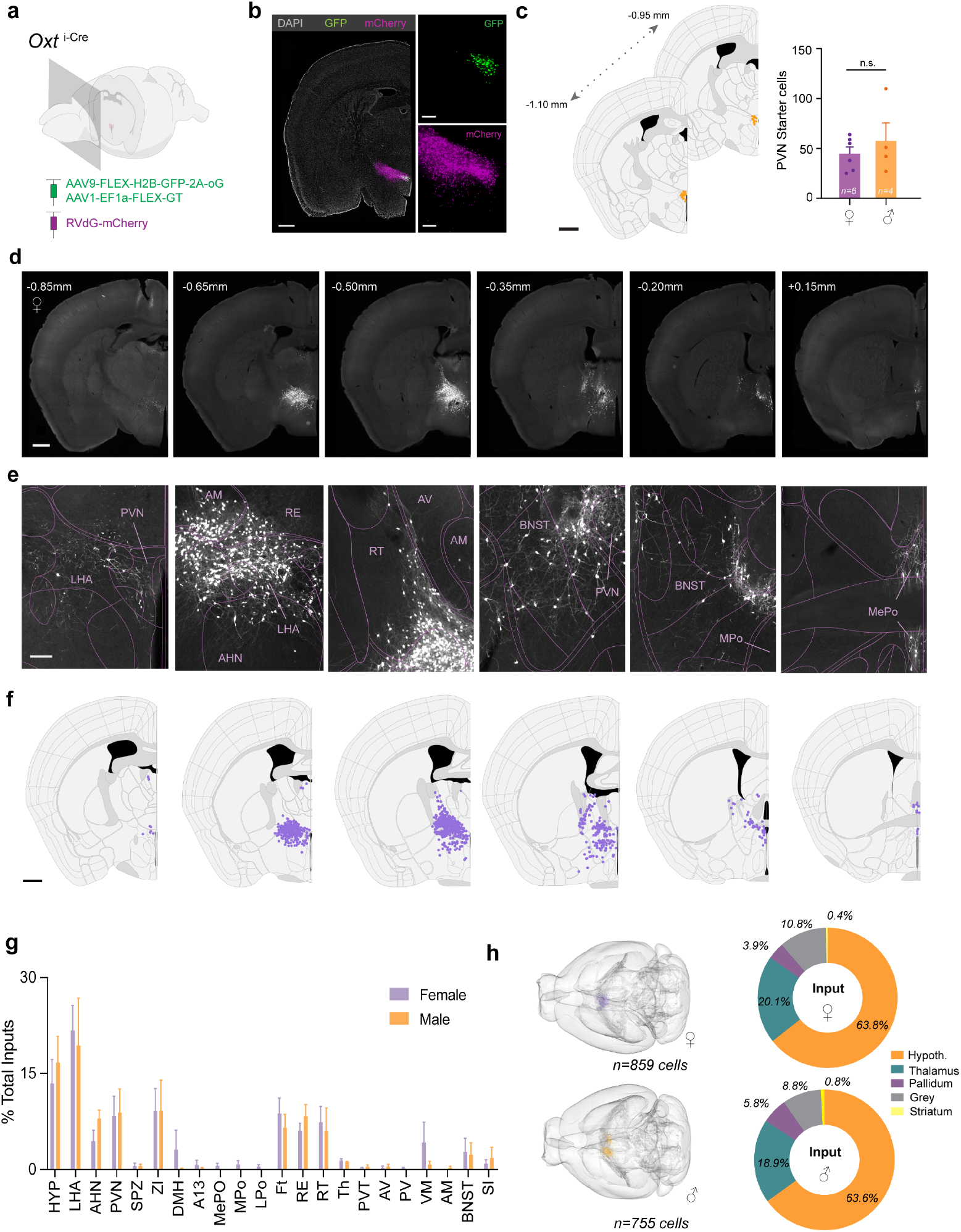
PVN Oxt+ inputs are largely restricted to surrounding hypothalamic and thalamic nuclei. a. Schematic of monosynaptic rabies tracing from PVN Oxt+ neurons. Coronal brain sections were collected at an interval of 180 µm. b. Representative image of starter cells in the PVN of male mouse. Starter cells are characterized as an overlap of GFP signal (green) and mCherry signal (magenta). Monosynaptic inputs are labeled with mCherry. Scale bar (left), 0.5 mm. Scale bars (right), 40 µm. c. *Left* Schematics of starter cell anterior-posterior spread in one male mouse brain. Starter cells are localized to PVN. Scale bar, 0.5 mm. *Right* Quantification of starter cells for female (n=6) and male (n=4) data sets. d. Representative raw images of retrogradely labeled inputs (mCherry+) to the PVN in a female mouse brain. Corresponding Bregma coordinates are depicted above each coronal slice. Inputs appear to be localized to hypothalamic and thalamic nuclei neighboring the PVN. Scale bar, 0.5 mm. e. Zoom in of retrogradely labeled input cells (mCherry+) from slices in panel d. Inputs are observed in hypothalamic, thalamic, and pallidal regions proximal to the PVN. Scale bar, 200 µm. f. Segmented monosynaptic inputs to Oxt+ neurons depicted in panel d and displayed on registered coronal slice from Allen brain reference atlas. Scale bar, 0.5 mm. g. Retrogradely labeled mCherry+ inputs by brain region, quantified as percentage of all retrogradely labeled cells. Regions depicted in the graph show on average about 1% or more total input. Male (n=4) and female (n=6) data do not show significant differences (adjusted p>0.9999, two-way ANOVA using Bonferroni post hocs for multiple comparisons). h. *Left* Schematic of all mCherry labeled cells projected into 3D space. In the representative female brain, 859 cells in total showed mCherry signal. Male brain shows a total of 755 mCherry+ cells. Both schematics demonstrate that Oxt inputs are highly proximal to PVN and do not extend to distal forebrain or hindbrain regions. *Right* Average total input of parent regions for female and male mice. The majority of inputs (>75%) are found in the hypothalamus (hypothal.)

Our data confirm that synaptic afferents to Oxt PVN neurons are primarily found in proximal hypothalamic and thalamic nuclei (Figure 2d-f). On average, 63.8% and 63.6% of inputs were from hypothalamic nuclei in female and male mice, respectively (Figure 2g-h). Our data show that the lateral hypothalamic area (LHA: 21.7±3.9% ♀, 19.4±7.4% ♂ of total input) and surrounding undefined hypothalamic regions (HYP: 13.4±3.7% ♀, 16.7±4.1% ♂) provide the majority of inputs to PVN Oxt neurons in both male and female mice (Figure 2f). As an integrative hub of peripheral sensory information, the LHA serves as a critical homeostatic regulator in the central nervous system. The complex cytoarchitecture of the LHA includes a diverse GABAergic neuronal population, providing a potential source of local inhibition to Oxt PVN neurons^37,38^. LHA inhibitory signaling may therefore function as an upstream regulator of Oxt synthesis and release from the paraventricular hypothalamic system. The zona incerta (ZI), anterior hypothalamic nucleus (AH), and the PVN itself also provide hypothalamic input to Oxt neurons, on a local circuit level, further supporting the possibility of local convergent input to regulate paraventricular neuronal firing (Fig 2f). Moreover, the Oxt system is capable of autoregulation due to the expression of cognate receptors within the PVN, pointing to multiple local hypothalamic mechanisms for control of Oxt release.

Surprisingly, we found that thalamic nuclei proximal to PVN account for a substantial fraction of projections to Oxt neurons. On average, 20.1% and 18.9% of inputs to Oxt neurons originated from the thalamus in female and male mice, respectively (Fig 2g). PVN Oxt neurons receive thalamic input primarily from the reticular nucleus (RT: 7.4±2.5% ♀, 6.0±3.6% ♂), nucleus of reuniens (RE: 6.1±1.1% ♀, 8.3±1.8% ♂), and ventral medial nucleus of the thalamus (VM: 4.2±3.2% ♀, 0.8±0.5% ♂) in both males and females (Figure 2g). Our data also determined that PVN Oxt neurons receive no cortical input (Figure 2-Figure Supplement 1d); thalamic relays may therefore serve as the key source of feedback information related to sensory processing of social behavior. Additionally, a small portion of PVN Oxt inputs were found in nuclei of the pallidum and fiber tract regions (Figure 2f-h). Pallidal regions projecting to Oxt neurons include the bed nucleus of the stria terminalis (BNST: 2.8±2.1% ♀, 2.3±1.9% ♂) and substantia innominata (SI: 0.9±0.6% ♀, 1.8±1.7% ♂), both previously described to mediate social and anxiety-related behaviors via oxytocinergic modulation. Our data support the existence of a reciprocally connected circuit that may be important for adapting or amplifying these behaviors. Finally, we found that inputs to PVN Oxt neurons are largely similar across males and females (Figure 2g, all adjusted p-values >0.9999, two-way ANOVA using Bonferroni method to correct for multiple comparisons). Together with our axonal mapping, the central oxytocin circuit appears to be sex-invariant.

### Brainwide axonal projections of paraventricular nucleus vasopressin neurons

Oxytocin and vasopressin circuits are thought to operate as an integrative system to regulate complex social behavior^4^. Dual release of both modulators and expression of their receptors have been reported in subcortical structures such as the amygdala, ventral striatum, and ventral tegmental area^3,13,39^. Yet, our understanding of whether these systems may work collaboratively or in a complementary manner is still limited. A quantitative full brain map of vasopressin (Avp) projections does not currently exist and could provide mechanistic insight to these possibilities. Avp output may target similar structures as the oxytocin system, or Avp neurons may tap into different nuclei, suggesting more complementary roles of the oxytocin and vasopressin circuits in the mammalian brain. To compare the output of the Avp and Oxt systems, we performed analogous anterograde tracing experiments in male and female mice, using Avp^*i-Cre*^ mice to target Avp+ neurons of the PVN^40^ (Figure 3a). Subjects exhibiting eYFP expression in the SON and the suprachiasmatic nucleus (SCN) were excluded from further analysis. Similar to the Oxt dataset, our quantitative analyses excluded cerebellar structures due to limited labeling in this region and therefore focused on cerebral cortex and subcortical regions. By employing the same analysis pipeline and experimental parameters (Figure 3c), we can more accurately compare Oxt and Avp projections, as well as determine whether the Avp system possesses sex-dependent features.

**Figure 3:**
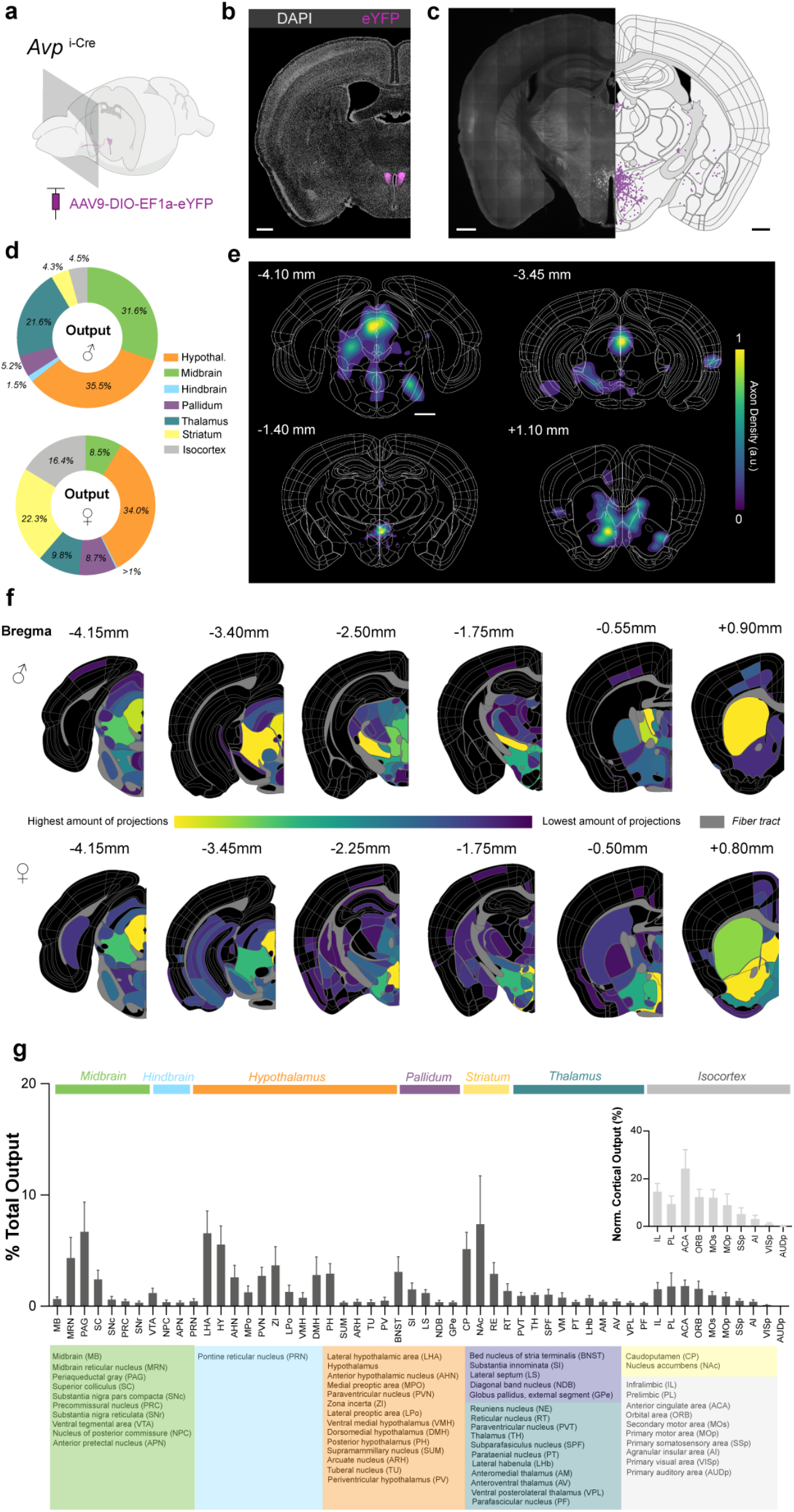
Quantitative anterograde map reveals broad, sex-dependent projections of Avp+ PVN neurons. a. Schematic of AAV9-DIO-eYFP injection into PVN of Avp^i-Cre^ mice. Coronal sections of the entire mouse brain were collected at an interval of 180 µm for projection mapping. b. Representative image of eYFP expression in Avp+ PVN neurons. Scale bar, 0.5 mm. *c. Left* Example raw image of Avp+ projections extending from the PVN to local hypothalamic regions. *Right* Registered coronal slice to Allen Brain atlas and segmented axonal projections of brain slice (*left*). Projections were identified using Sobel edge detector algorithm and quantified as pixel counts. d. Comparison of average total axonal output of Avp+ neurons across male (n=2) and female (n=3) mice in parent regions of Allen brain atlas. These regions include the hypothalamus (hypothal.), midbrain, hindbrain, pallidum, striatum, and isocortex. Avp anterograde tracing largely revealed differences in projection patterns across male and female mice. e. Projection density heatmaps of representative female mouse coronal sections. Density contour maps are generated using two-dimensional kernel density estimation across a square grid. Regions with greatest density of projections are depicted in yellow, while regions containing few projections are depicted in purple. Bregma coordinates are depicted above each registered brain slice. Scale bar, 1 mm. f. Coronal heat maps of one male and one female mouse. Each nucleus displays a color corresponding to normalized amount of projections. Nuclei with highest number of projections are depicted in yellow, while areas with little to no projections are displayed in purple or black. Male and female mice show distinct differences in projection intensity across the brain, particularly in striatal and midbrain nuclei. g. Brain nuclei showing Avp+ projections in male and female mice (n=7 total), quantified as % of total quantified projections in the brain. Regions depicted in graph show >0.2% of total projections. For a list of all quantified regions, see Table 2. Inset, cortical regions normalized to total isocortical pixel count. Brain region acronyms with corresponding parent regions are depicted below.

Avp projections were found dispersed throughout the brain, including distal cortical and hindbrain regions (Figure 3d, Table 2). Avp projection patterns appeared to vary more strongly by sex, compared to the Oxt system. Male mice showed greater eYFP cell labeling in extra-paraventricular nuclei, located adjacent to the PVN (Figure 3-Figure Supplement 1a-c). Both vasopressin and oxytocin neurons have been reported to exist in lower abundance in nuclei outside the PVN^41–43^. The additional labeling of neurons adjacent to the PVN in male mice may represent a sex-variant feature of the vasopressinergic system that could potentially underlie the differences in projection patterns seen across male and female mice, but further research is needed to evaluate this possibility and the function of these extra-PVN neurons.

**TABLE 2.**
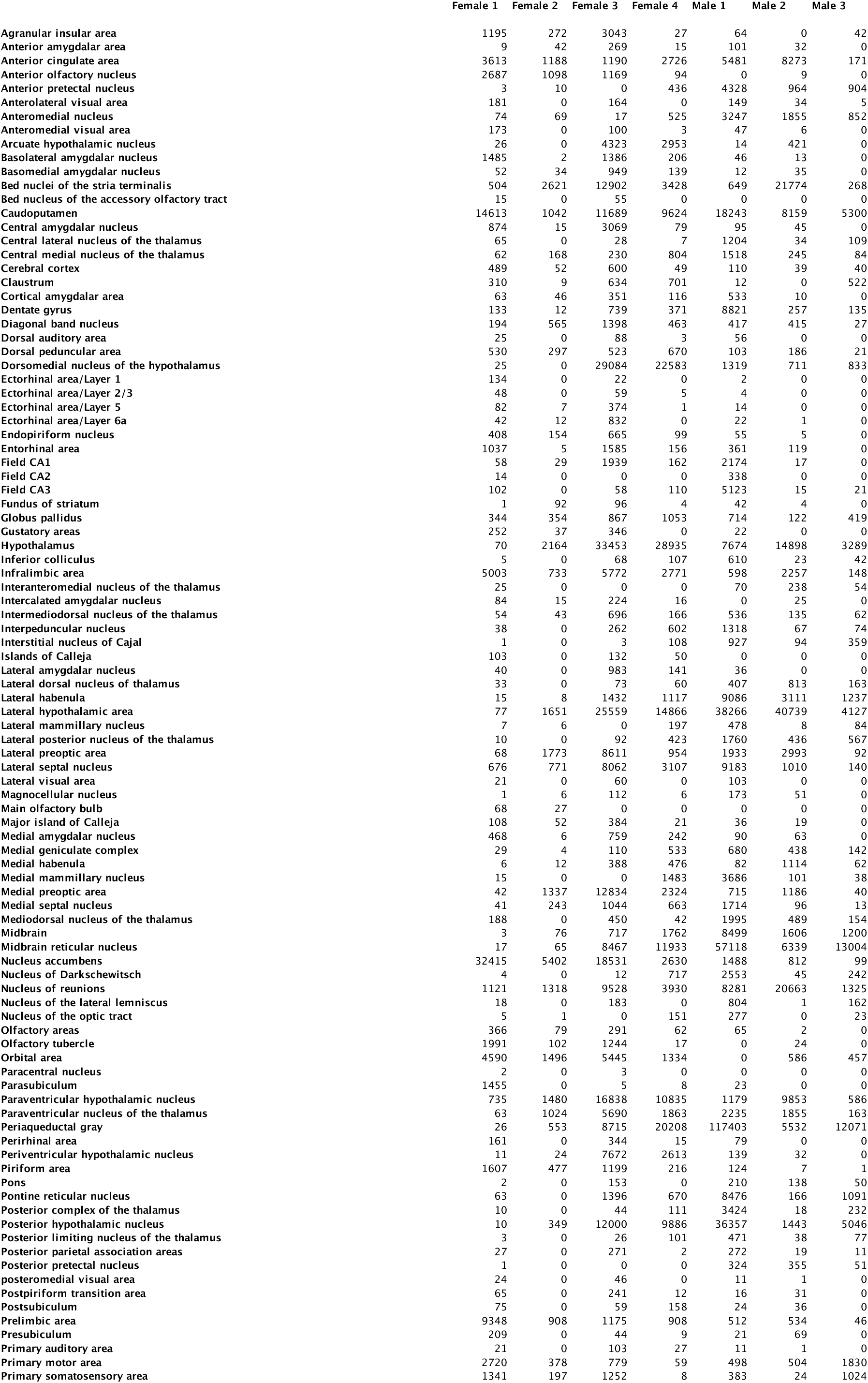
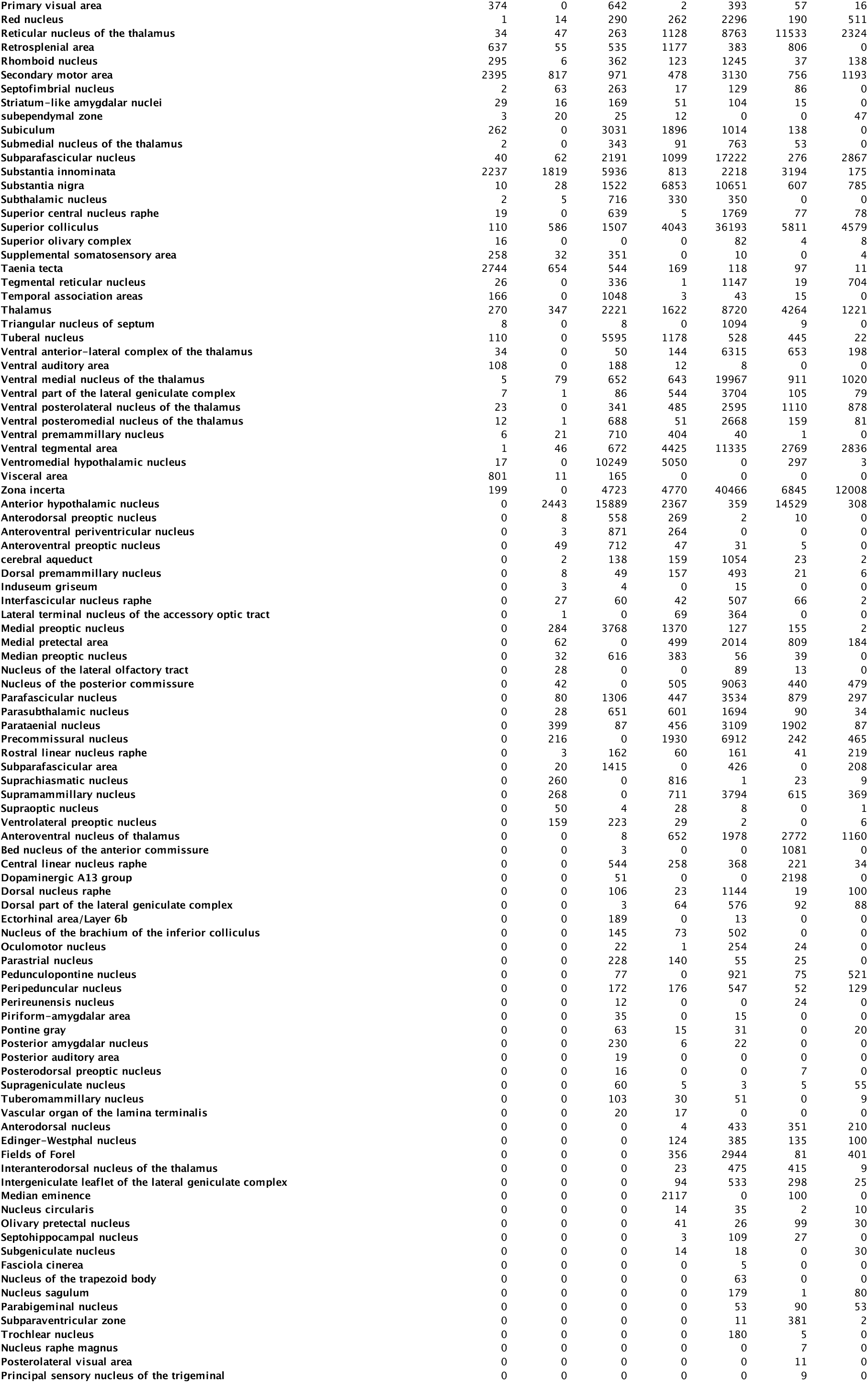
Total vasopressin projection pixel counts per sample (related to Figure 3)

In females, Avp output targeted more forebrain structures, primarily striatal and local hypothalamic nuclei (Figure 3d, Figure 3-Figure Supplement 2a-c). On average, 36.1% of total output targeted hypothalamic nuclei and 33.7% of all output extended to striatal regions in female mice (Figure 3d). By comparison, male mice showed less dense axonal labeling in striatal regions, accounting for an average of 4.6% total output. Avp projections to hypothalamic areas include the zona incerta (ZI: 0.9±0.5% ♀, 7.3±2.8% ♂, adjusted p-value=0.04, two-way ANOVA using Bonferroni correction for multiple comparisons), lateral hypothalamic area (LHA: 4.5±1.6% ♀, 9.3±4.1% ♂), anterior hypothalamic nucleus (AHN: 2.9±1.4% ♀, 2.2±2.0% ♂), and dorsal medial hypothalamic nucleus (DMH: 4.6±2.6% ♀, 0.5±0.2% ♂). Compared to Oxt projections, Avp+ neurons in females target more ventral striatal structures including the nucleus accumbens (NAc: 12.7±6.6% ♀, 0.2±0.07% ♂), aligning with the known role of Avp signaling in the formation of affiliative bonds^13^ (Figure 3e-f). This axonal pattern was statistically different than that observed in males (p<0.0001, two-way ANOVA using Bonferroni correction for multiple comparisons). Interestingly, striatal projections in males were largely localized to dorsal striatum, termed caudate putamen (CP: 6.0±2.7% ♀, 3.9±0.7% ♂) in Allen brain atlas ontology. Additionally, Avp isocortical targets contributed a much larger proportion of total axonal output in female mice (Figure 3d-f). Avp fibers were observed in medial prefrontal cortex nuclei—infralimbic, prelimbic, and anterior cingulate cortices— as well as in the orbital cortex.

Our anterograde tracing also determined that male mice exhibited distinct Avp+ projection patterns in midbrain and hindbrain regions compared to females. In males, most PVN Avp+ output was directed to hypothalamic and midbrain structures, more closely modeling Oxt projections (Figure 3d, Figure 3-Figure Supplement 1). On average, 48.8% total Avp output was directed towards midbrain nuclei and 27.7% of output targeted hypothalamic nuclei (Figure 3d). Both the periaqueductal gray (PAG: 3.2±2.0% ♀, 11.3±4.9% ♂, adjusted p=0.001) and midbrain reticular nucleus (MRN: 1.5±1.0% ♀, 8.2±3.1% ♂, adjusted p=0.02) showed statistically significant differences in males and females. Avp projections to the reward related ventral tegmental area (VTA: 0.6±0.5% ♀, 2.0±0.5% ♂) and substantia nigra pars compacta (SNc: 0.6±0.5% ♀, 0.6±0.2% ♂) were far less dense compared to projections from the Oxt system, a pattern consistent in females and males (Figure 3f-g, Figure 3-Figure Supplement 1 and 2). In addition, males exhibited more Avp+ projections in thalamic structures, accounting for 13.6% of total output. Nuclei with PVN Avp+ fibers included the lateral habenula (LHb: 0.2±0.1% ♀, 1.4±0.1% ♂), nucleus of reuniens (RE: 2.2±0.5% ♀, 3.8±2.5% ♂), and reticular nucleus (RT: 0.2±0.1% ♀, 2.9±1.0% ♂). Together, our anterograde tracing results suggest that vasopressinergic output may be sexually dimorphic in mice. Our data align with a recent study looking at immunoreactive Avp and Oxt fibers in socially relevant nuclei of the rat brain^44^. In both hypothalamic and striatal regions, the authors found sex differences in Avp projections, but not in Oxt axons. Additionally, our brain-wide mapping revealed that the distribution of Oxt and Avp axonal projections is distinct, suggesting that the operations of these two modulatory systems may independently regulate behavior and homeostatic mechanisms.

### Retrograde input mapping of paraventricular nucleus vasopressin neurons

Previous studies using non-genetically targeted retrograde tracers have found presynaptic inputs to PVN neurons from distal structures, including amygdalar and septal nuclei^45^. Considering the heterogeneity of cell types in the PVN, these studies provided a limited knowledge of synaptic afferents specific to vasopressin neurons. A more recent study using genetically-targeted monosynaptic rabies tracing compared input to Avp+ PVN and SON neurons and found synaptic inputs ranging from limbic to midbrain structures^46^. Yet, this tracing study does not compare across sexes or allow for alignment to the oxytocin system, under the same experimental conditions. We performed quantitative, brainwide input mapping of PVN Avp+ neurons to determine how Oxt and Avp synaptic input may differ and whether Avp+ input may be sex dependent. These experiments and analyses are analogous to those performed for Oxt neurons, allowing for a more controlled comparison of the two modulatory systems (Figure 4a-c, Figure 4-Figure Supplement 1).

**Figure 4:**
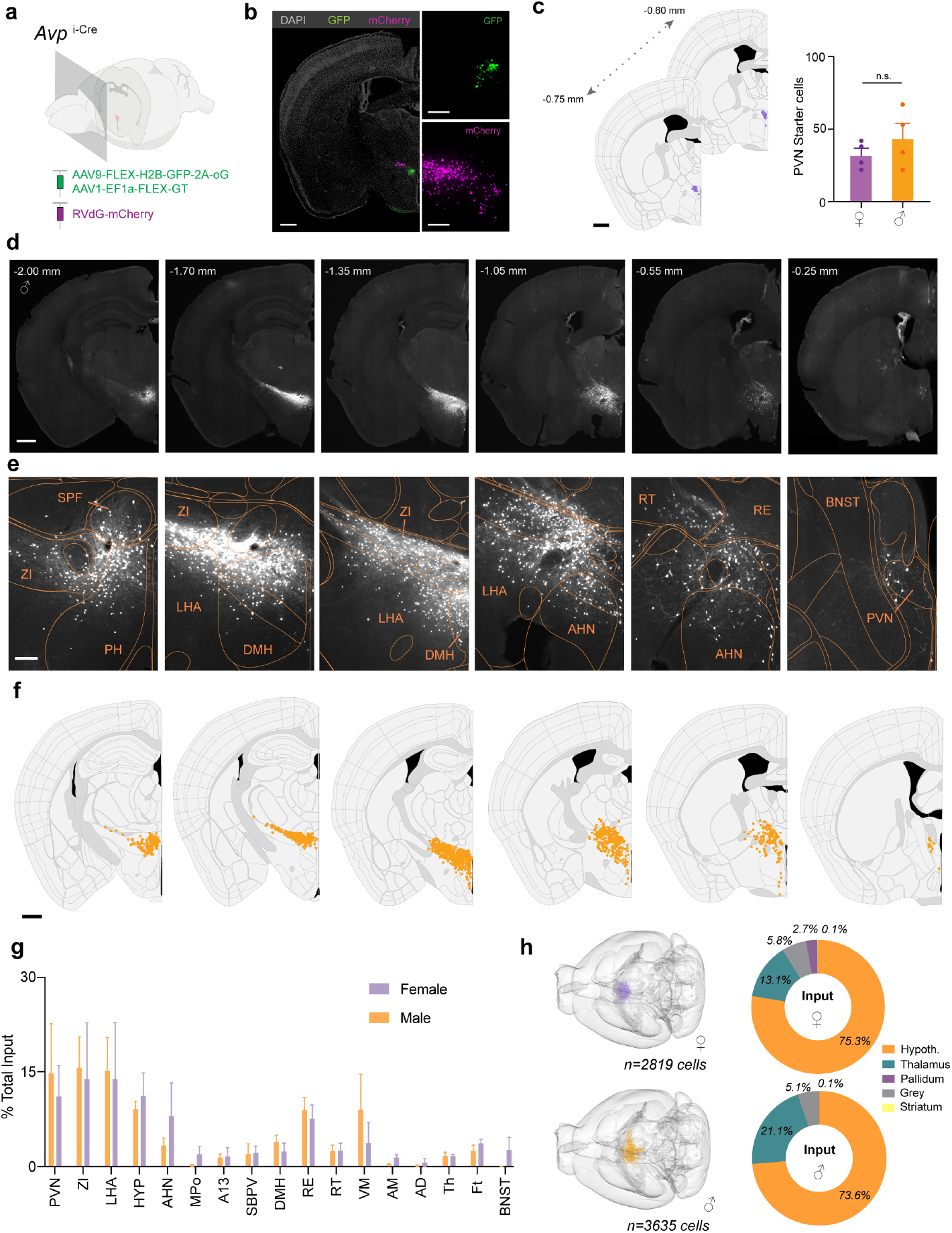
PVN Avp+ inputs are observed in surrounding hypothalamic and thalamic nuclei. a. Schematic of monosynaptic rabies tracing from PVN Avp+ neurons. Coronal brain sections were collected at an interval of 180 µm. b. Representative image of starter cells in the PVN of female mouse. Starter cells are characterized as an overlap of GFP signal (green) and mCherry signal (magenta). Monosynaptic inputs are labeled with mCherry. Scale bar (left), 0.5 mm. Scale bars (right), 40 µm. c. *Left* Schematics of starter cell anterior-posterior spread in one male mouse brain. Starter cells are localized to PVN. Scale bar, 0.5 mm. *Right* Quantification of starter cells for female (n=4) and male (n=4) data sets. d. Representative raw images of retrogradely labeled inputs (mCherry+) to PVN Avp+ neurons in a male mouse brain. Corresponding Bregma coordinates are depicted above each coronal slice. Inputs appear to be localized to hypothalamic and thalamic nuclei neighboring the PVN. Scale bar, 0.5 mm. e. Close up images of labeled input cells (mCherry+) of slices shown in panel d. Scale bar, 200 µm. f. Segmented monosynaptic inputs to Oxt+ neurons depicted in panel d and displayed on a registered coronal slice from Allen brain atlas. Scale bar, 0.5 mm. g. Retrogradely labeled mCherry+ inputs by brain region, quantified as percentage of all retrogradely labeled cells. Regions depicted in the graph show more than 1% total input. Male (n=4) and female (n=4) data do not show significant differences (adjusted p>0.9999, two-way ANOVA using Bonferroni post hocs for multiple comparisons). h. *Left* Schematic of all mCherry labeled cells projected into 3D space. In the representative female brain, 859 cells in total showed mCherry signal. Male brain shows a total of 755 mCherry+ cells. In both males and females, Avp inputs are highly proximal to PVN and do not extend to distal forebrain or hindbrain regions. *Right* Average total input of parent regions for female and male mice. The majority of inputs (>70%) are found in the hypothalamus (hypoth.) and thalamus.

The Avp+ neuron input map showed many similarities to PVN Oxt+ neurons. Synaptic afferents to PVN Avp neurons were largely restricted to neighboring hypothalamic nuclei, and the inputs appeared similar across male and female mice (Figure 4d-f). On average, 75.3% and 73.6% of inputs were from hypothalamic nuclei in female and male mice, respectively (Figure 4h). Hypothalamic regions providing substantial input to Avp neurons in both males and females included the lateral hypothalamic area (LHA: 13.9±8.9% total input ♀, 15.2±5.4% ♂), zona incerta (ZI: 13.9±8.9% ♀, 15.6±4.9% ♂), and anterior hypothalamic nucleus (AHN: 8.0±5.3% ♀, 3.4±1.1% ♂). A large fraction of inputs was also found in undefined hypothalamic nuclei in the Allen reference atlas (HYP), contributing 11.2±3.6% input in females and 9.1±1.2% of input in males. As observed in the oxytocinergic system, we also found PVN input to Avp+ neurons (Figure 4e-f), suggesting a potential conserved feedback mechanism to control Avp and Oxt release.

Beyond local hypothalamic nuclei, our data found inputs to Avp+ neurons from thalamic nuclei (Figure 4g-h). On average, thalamic nuclei contributed 13.1% of total input in females and 21.1% in males (Figure 4g). Thalamic nuclei sending synaptic input to Avp+ and Oxt+ PVN neurons were the same, potentially representing another common regulatory mechanism to control the release of each neuropeptide. Shared thalamic inputs include: the nucleus of reuniens (RE: 7.6±2.1% ♀, 9.0±1.9% ♂), ventromedial thalamic nucleus (VM: 3.7±3.2% ♀, 9.0±5.6% ♂), and reticular nucleus (RT: 2.5±1.2% ♀, 2.5±0.9% ♂). The remaining inputs to Avp+ neurons were largely found in fiber tract regions—contributing approximately 5% of total input in males and females— and in pallidal nuclei (Figure 4f-h). Interestingly, pallidal input showed a slight, but not statistically significant difference across male and female mice. Pallidal input made up 2.7% of total input in females, largely from the bed nucleus of the stria terminalis (BNST: 2.6±2.0%), while males exhibited nearly no input from this area (0.01±0.01%, adjusted p>0.99, two-way ANOVA Bonferroni correction for multiple comparisons). In combination with our axonal mapping data, the BNST emerges as a candidate sexually-dimorphic circuit for vasopressin neuromodulation. Overall, our Avp+ PVN input map revealed many similarities to Oxt+ neurons and demonstrated that synaptic afferents to these peptidergic populations are largely sex-invariant.

## Discussion

We used genetic viral tracing methods to map the input-out architecture of the paraventricular oxytocin and vasopressin systems in mice. Using modified open-source software^22^ and user-friendly accessible 2D coronal section analysis, our results present a unified resource to delineate and compare the basic structure of hypothalamic oxytocin and vasopressin circuitry. Our analyses also include systematic comparisons across male and females to determine whether sex-variant features exist at the level of synaptic afferents or efferents. These data help answer whether oxytocin and vasopressin exert their sexually dimorphic behavioral output via central or peripheral mechanisms. We find that both nonapeptide circuits are largely sex-conserved, with vasopressin neurons exhibiting some key differences in axonal output between male and female mice. The use of complementary methods side by side allows for a more controlled comparison of both neuropeptidergic systems to determine the extent to which oxytocin and vasopressin may operate in a collaborative or complementary manner. Our retrograde input maps found numerous similarities in both systems, suggesting conserved regulatory mechanisms for control of neuronal activity in the two populations.

Neurohypophyseal peptides are linked to sex-specific behaviors^6,30,47^, yet it is unclear if differences in central anatomic circuitry subserve sex-dependent physiological functions. Together, our data suggest that the central oxytocin system is largely sex-invariant. Consistent in both males and females, most presynaptic inputs to paraventricular oxytocin neurons are restricted to surrounding hypothalamic nuclei. We did not find evidence of long-range synaptic afferents from cortical, midbrain, or hindbrain regions. Instead, our retrograde data suggest that oxytocin release is potentially regulated by a localized hypothalamic circuit. Previous electrophysiological mapping of inputs to paraventricular neurons supports the existence of numerous proximal hypothalamic contacts, including from local GABAergic neurons^32–34,48^. Considering the heterogeneity of cell types in the lateral hypothalamus, more work is needed to understand what hypothalamic cells project to oxytocin neurons and how their activation might alter or regulate oxytocin neuronal firing^49^. One possibility supported by lesion and stimulation studies is that more proximal regions such as limbic nuclei (*e*.*g*., septum, amygdala, subiculum) can control oxytocin release through feed-forward inhibition from local hypothalamic GABAergic neurons^48,50^. This is aligned with anatomical tracing of fibers from these areas that find terminals surrounding, but not within the PVN^33^. The understanding of PVN and LHA interactions is complicated by the close proximity of these two brain regions, which makes interpreting neuroanatomical results more challenging compared to distal regions (*e*.*g*., Fig 2d-e, Fig S5). Fine-scale understanding of neuroanatomy in such closely located hypothalamic structures may be resolved with extremely sparse labeling methods or orthogonal genetic targeting approaches.

In addition to the hypothalamus, we find that thalamic nuclei provide substantial input to oxytocin neurons. Oxytocin neuron activity can change in response to diverse stimuli, including pup distress calls, social interaction, and social touch^18,30,51^. Because we did not find evidence of cortical inputs unlike previous studies, a substantial proportion of these peripheral sensory cues may be relayed through thalamic structures in mice^36^. Oxytocin release has been reported in nearly all sensory-related cortical regions, yet far less is known about how sensory cues are relayed and processed to regulate oxytocin release^17^. Thalamic inputs found in our retrograde tracing data, such as the nucleus of reuniens and reticular nucleus, emerge as potential candidates for thalamic processing of socially relevant sensory information. Processing of social cues is likely sex-dependent—somatosensory cues for lactation, pup distress calls, and nursing are highly salient for female rodents, but not for males. Our retrograde data provides the first evidence that central components of Oxt circuit architecture likely do not underlie these differences, suggesting that they may be linked to sex-dependent distributions of receptors or peripheral mechanisms. Afferents to paraventricular vasopressin neurons exhibited numerous parallels to the oxytocin system, with prominent inputs from the nucleus of reuniens, reticular nucleus, and ventromedial thalamic nucleus. These regions may therefore serve as more generalized processing centers of salient social cues that can modulate the gain of input activity from distal sensory cortices. The vasopressin- and oxytocinergic systems may exert unilateral modulatory control over sensory-related cortical regions, but more complex integration of peripheral sensory information—related to emotional and social processing—may be necessary to provide feedback and ultimately gate peptidergic release.

Our quantitative anterograde mapping is the first brain-wide, comparative study of the output patterns of paraventricular vasopressin and oxytocin neurons. We find that the axonal projection patterns of each system exhibit slight variations, with oxytocin fibers more predominantly found in midbrain structures and vasopressin fibers found in more forebrain regions, including greater contributions to cortical and striatal nuclei. Only vasopressinergic projections showed substantive sex-variant features, potentially driven by the contribution of a small population of extra-paraventricular vasopressin+ neurons in male mice. Our brain-wide anterograde maps also highlight areas of oxytocin and vasopressin modulation that are currently understudied. This includes robust innervation of the midbrain reticular nucleus, pontine reticular nucleus, and periaqueductal gray where the modulatory effects of oxytocin and vasopressin release can tune neuronal activity to regulate motivated behaviors. Oxytocin and vasopressin are often regarded as “sister” peptides that form a highly integrative, cooperative system to gate social behavior^1^. Despite their similarities, each peptide controls different physiological functions. The divergent axonal patterns in our anterograde maps provide a circuit basis for how oxytocin and vasopressin may operate in a complementary manner to control different homeostatic or social behaviors.

Together, our data provide insight into the circuit architecture of neurohypophyseal peptides. Oxytocin, unlike vasopressin, appears more sex-conserved in its input-output architecture. Minimal sex-variant patterns of inputs to and outputs from paraventricular oxytocin neurons suggests that central circuit architecture likely does not underlie its sex-specific functions. Instead, differences in receptor distributions across males and females may provide a method of differential modulation, which has been previously reported in rodent brains^20,47,52^. Alternatively or in addition, peripheral mechanisms and non-canonical pathway interactions between oxytocin and other neuropeptides may provide new avenues to understand how oxytocin mediates different behaviors. This side-by-side comparison of vasopressin and oxytocin central circuit organization provides a road map to study how these two pleiotropic peptides work cooperatively and independently to control a wide variety of animal behaviors.

## Materials and Methods

### Mouse strains and genotyping

Animals were handled according to protocols approved by the Northwestern University Animal Care and Use Committee. Male and female mice were used for all experiments, and all experiments were performed on young adult (postnatal day >35) mice. C57BL/6 mice used for breeding were acquired from Charles River (Wilmington, MA); other mouse lines were acquired from the Jackson Laboratory (Bar Harbor, ME). B6.129S-*Oxt* ^tm1.1(cre)Dolsn^/J mice (#024234, referred to as *Oxt* ^i-Cre^), which express the enzyme Cre recombinase under control of the oxytocin promoter^24^, were used to target oxytocinergic neurons. B6.Cg-*AVP* ^tm1.1(cre)Hze^ mice (#023530, referred to as *Avp* ^i-Cre^), which express the enzyme Cre recombinase under control of the vasopressin promoter, were used to target vasopressin neurons^40^. Mice heterozygous for Cre were used for all retrograde and anterograde tracing experiments. All genotyping primers were based on standard protocols available on the Jackson Lab website.

### Stereotaxic intracranial injections

Young adult mice (>P35) of both sexes were anesthetized with 1.5-2% isofluorane and positioned on a small stereotaxic frame (David Kopf Instruments, Tujunga, CA) so the skull was lying flat with the vertical positions of lambda and bregma within 0.1 mm of each other. Wiretrol II pipettes (Drummond Scientific Company) pulled on a P-1000 Flaming/Brown micropipette puller (Sutter Instruments) were used for all injections. Both retrograde and anterograde tracing experiments used unilateral injections at two different depths below the pial surface. To target the PVN, the pipette was placed 0.4 mm lateral to the midline (+0.4mm M/L), 0.5 mm posterior to bregma (−0.4 mm A/P), and both 4.4 and 4.7 mm ventral to the pial surface (−4.4 and -4.7 mm D/V). Virus was injected using a microsyringe pump (World Precision Instruments); >5 minutes elapsed before the pipette was moved from the ventral injection site to the more dorsal injection site, and 10 minutes elapsed before the pipette was slowly retracted from the brain following the second injection. Mice were given ketoprofen (5 mg/kg) for analgesia and recovered on a heating pad before being returned to their home cage. Two additional doses of ketoprofen were administered 24 and 48 hours post-operation and mice were closely monitored by research staff for signs of distress or pain following surgical procedures.

For trans-synaptic retrograde tracing, 200 nl of a 1:1 mixture of AAV9-FLEX-H2B-GFP-2A-G (8.6×10^12^ GC/ml, Salk Institute, La Jolla, CA) and AAV1-EF1a-FLEX-GT (8.6×10^12^ GC/ml, Salk Institute) were unilaterally injected into the PVN (0.4 mm posterior to bregma, 0.4 mm lateral, and 4.4 mm and 4.7 mm below the pia) of *Oxt* ^i-Cre^ and Avp ^i-Cre^ mice at a rate of 100 nl/min, using the procedure described above. Three weeks later, 400 nl of EnvA G-deleted Rabies-mCherry was injected into the same area of the PVN with 200 nl injected at each site at a rate of 100 nl/min. After recovery, mice were housed in a biosafety level 2 (BSL2) facility for 1 week before euthanasia.

For anterograde tracing, a total of 300-400 nl of AAV9-EF1a-DIO-eYFP-WPRE (5.55 × 10^12^ gc/ml, University of Pennsylvania Vector Core, Philadephia, PA) was unilaterally injected at -4.4 and -4.7 D/V at a rate of 100 nl/min in Oxt ^i-Cre^ or Avp ^i-Cre^ mice (150-200 nl per injection site). Mice were sacrificed and fixed 3-4 weeks after virus injection.

### Tissue processing and immunohistochemistry

Mice were anesthetized with isoflurane and perfused transcardially with 4% paraformaldehyde (PFA) (Electron Microscopy Sciences) in phosphate buffered saline (PBS). Brains were extracted and stored in 4% PFA 1-2 days post extraction, then stored in PBS. Coronal brain slices were made at 60 µm thickness for both fluorescent projection mapping and retrograde tracing. All slices were made using a Leica VT1000 S vibratome (Leica Biosystems). One in every three sections (180 µm sample frequency) was used for imaging, expression validation, and when applicable, further immunostaining. Subjects that showed expression in the supraoptic nucleus (SON) were excluded from analyses.

For immunohistochemistry, sections were pretreated in 0.2% triton-X100 for an hour at RT, then blocked in 10% bovine serum albumin (BSA, Sigma-Aldrich, ST Louis, MO):PBS with 0.05% triton-X100 for two hours at RT, and they were then incubated overnight at 4°C with primary antibody solution in PBS with 0.2% triton-X 100. On the following day, tissue was rinsed in PBS, reacted with secondary antibody for 2 hours at RT, rinsed, mounted onto Superfrost Plus slides (ThermoFisher Scientific, Waltham, MA), dried and coverslipped under glycerol:TBS (3:1) with Hoechst 33342 (1:1000; ThermoFisher Scientific). Primary antibodies with the following dilutions used in this study include: rabbit anti-oxytocin (1:1000; T-4084, Peninsula Laboratories International, Inc., San Carlos, CA), rabbit anti-vasopressin (1:1000; AB1565 MilliporeSigma, Burlington, MA), chicken anti-GFP (1:2000, ab13970, Abcam, Cambridge, UK), rabbit anti-RFP (1:1000, 600-401-379, Rockland Immunochemicals, 600-401-379, Pottstown, PA). Alexa Fluor 488- and 647-conjugated secondary antibodies against rabbit or chicken (Life Technologies, Carlsbad, CA) were diluted 1:500.

### Anatomical imaging and registration

Whole brain sections with viral fluorescent expression or immunolabeling were imaged on an Olympus VS110 imaging system at 10x (Olympus Scientific Solutions Americas, Waltham, MA). For anterograde mapping experiments, samples were not immunoenhanced, and eYFP viral expression was used for projection quantification. For retrograde mapping, mCherry and GFP signals were immunoenhanced for starter cell expression validation; however, non-immunoehanced mCherry signal was used for quantification of monosynaptic input neurons. All images were converted from Olympus .vsi file format to .tiff (16-bit images) for analysis using an automated batch-processing script in FIJI (ImageJ) to ensure standardized image format for quantification.

Quantitative two-dimensional projection and retrograde input mapping was performed using the the Allen Mouse Brain Reference Atlas and an adapted version of the open-source software, WholeBrain (Fürth et al., 2017), a mouse brain registration package in R. Briefly, the original open-source R package was designed to spatially map neuroanatomical data at the cellular-level to the Allen standardized mouse atlas^23^. Our modified analyses implemented a Sobel edge-detection method to identify axonal processes and multi-otsu thresholding to segment somatic signal. For registration with the WholeBrain package, autofluorescence of brain sections is used to register coronal slices to Allen reference atlas and segmented axonal and somatic signals are transformed to the reference brain section. Segmented axons were quantified as pixel counts for each registered brain region, and each sample was normalized to total axonal content to account for potential variability in injection site locations or viral expression across samples. Cerebellar and hindbrain coronal slices were excluded from quantification due to low number of eYFP+ projections (see Figure 1-Figure Supplement 1).

For retrograde tracing quantification, inputs (mCherry+ cells) were quantified as total percentage of all retrogradely labeled cells to normalize for differences in number of starter cells across samples. Starter cells were defined as cells co-labeled with mCherry and GFP signal; and these cells were excluded from final quantification. To locate starter cells, coronal slices containing GFP signal were registered and segmented using the same parameters established for retrograde tracing of mCherry+ slices. Centroid x-y coordinates and area of GFP+ cells were recorded, and the Euclidean distance was calculated between mCherry+ cells in the same slice. Signals were considered to overlap if the Euclidean distance was < 5 µm, and the cell was categorized as a starter cell.

Axonal segmentation was performed using the ‘OpenImageR’ package. Coronal axon density contours and heatmaps were generated in R using ‘tidyverse’ and ‘ggplot2’ packages with the “viridis” package for visualization. For density contours depicted in Figure 1e, Figure 3e, and Figure 1-Figure Supplement 2-3 & Figure 3-Figure Supplement 1-2, we used the ‘geom_density_2d’ function that estimates two-dimensional kernel density with an axis-aligned bivariate normal kernel, evaluated on a square grid. The results are displayed as density contours that are overlayed with outlines of the registered coronal brain slice from Allen atlas. Coronal heatmaps in Figure 1d-e and Figure 3d-e were also generated using the ‘tidyverse’ and ‘ggplot2’ packages. Segmented axons for each subregion were counted and normalized to total pixel count for the entire sample. The square root of the normalized pixel count was taken for each region and then assigned a color fill from the ‘viridis’ color scheme to represent pixel count or intensity. This color fill was then plotted with the registered Allen brain atlas coronal slice using outlines from the ‘WholeBrain’ package. For the matrix heatmap plots depicted in Figure 1-Figure Supplement 4 and Figure 3-Figure Supplement 3, the square root of normalized pixel counts for each region was also used to assign a color from the ‘virdis’ color scheme. Rows were clustered using the ‘dendrogram’ function of the ‘tidyverse’ package.

### Quantification and statistical analyses

All 2D brain registration, segmentation, and data visualization were performed in R, version 3.6.3 for Mac OSX. Quantification and statistical analysis were performed in Matlab (2020a, Mathworks) and Prism 8.4 (GraphPad Software). All statistical tests were two-sided. Statistical details of experiments can be found in the results section and figure legends. For comparison of male and female data, two-way ANOVAs were performed using Bonferroni correction for multiple comparisons. Statistical significance was set as alpha=0.05. Comparisons between unpaired data were performed using two-tailed Mann-Whitney tests. Summary data in all main figures are reported as mean ± SEM. All figures were compiled in Adobe Illustrator (2021).

---

**Figure 1—Figure Supplement 1:**
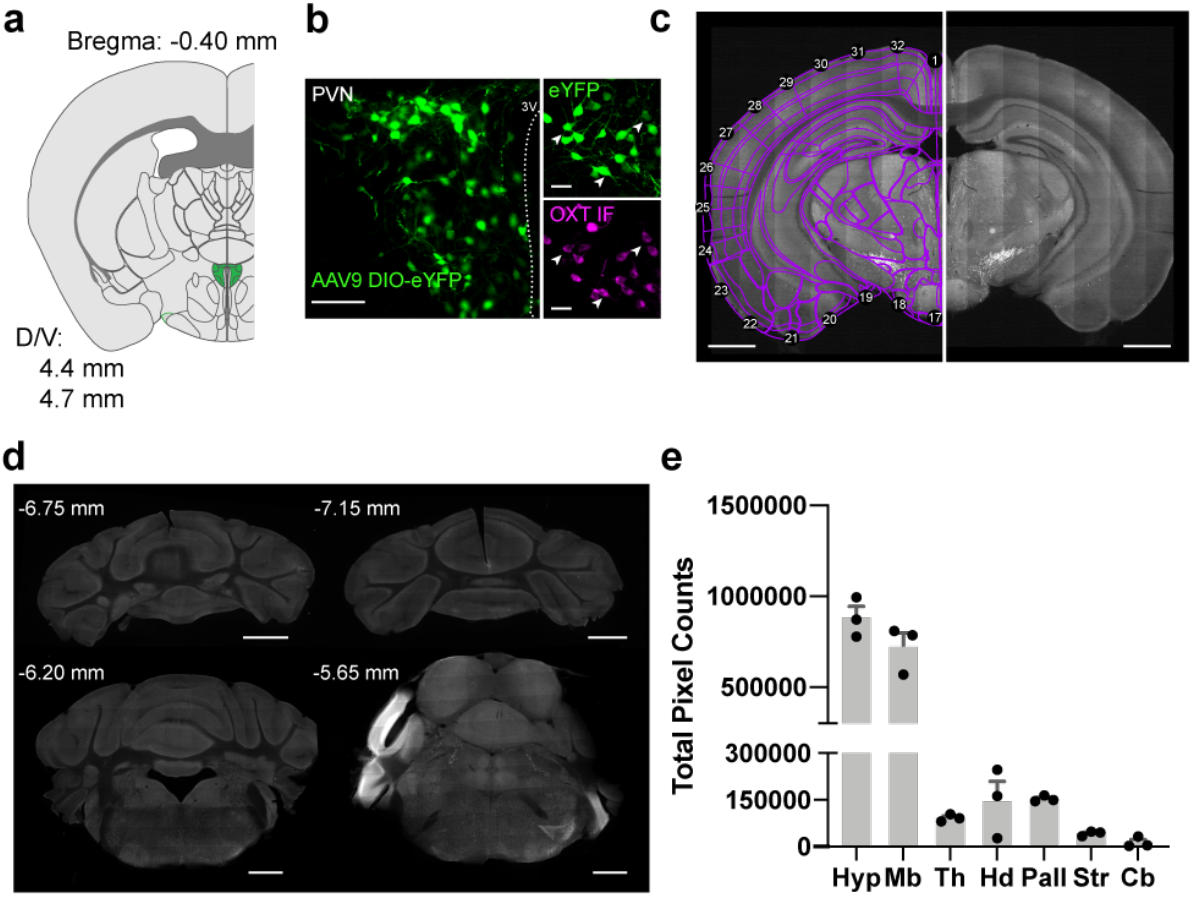
Workflow for registration and quantification of brainwide Oxt+ axons. a. Bregma coordinates for targeted injections to the PVN in adult male and female mice. A total of 300-400 nl of a Cre-dependent eYFP virus was injected at 4.4 and 4.7 mm below the pia (150-200 nl per injection site). Slice represents PVN region where densest eYFP expression was observed in all samples (Bregma: -0.70mm). b. Expression of AAV9-DIO-eYFP was restricted to Oxt+ neurons in the PVN, Scale bar 100 µm. *Inset* Confocal images of neurons with eYFP expression and immunolabeling for Oxt. White arrows depict cluster of neurons co-labeled with eYFP and Oxt antibody, Scale bar 40 µm. c. Example raw image of Oxt projections and registration using WholeBrain software package in R. A series of correspondence points (labeled numbers) are used to align coronal slices of the Allen brain atlas to 2D coronal slices containing eYFP expression. Scale bar, 1 mm. d. Representative cerebellar cortex and hindbrain images from one male and one female mouse brain exhibiting little eYFP anterograde signal. Corresponding Bregma coordinates are depicted above each image. Scale bar, 1 mm. e. Quantification of total pixel counts for three brains (n=1 male, 2 female) across parent regions of the mouse brain: the hypothalamus (Hyp), thalamus (Th), midbrain (Md), hindbrain (Hd), pallidum (Pall), striatum (Str), and cerebellum (Cb). Cerebellum shows limited pixel intensity in both male and female samples. PVN projections appear to be largely localized to cerebral cortex structures.

**Figure 1—Figure Supplement 2:**
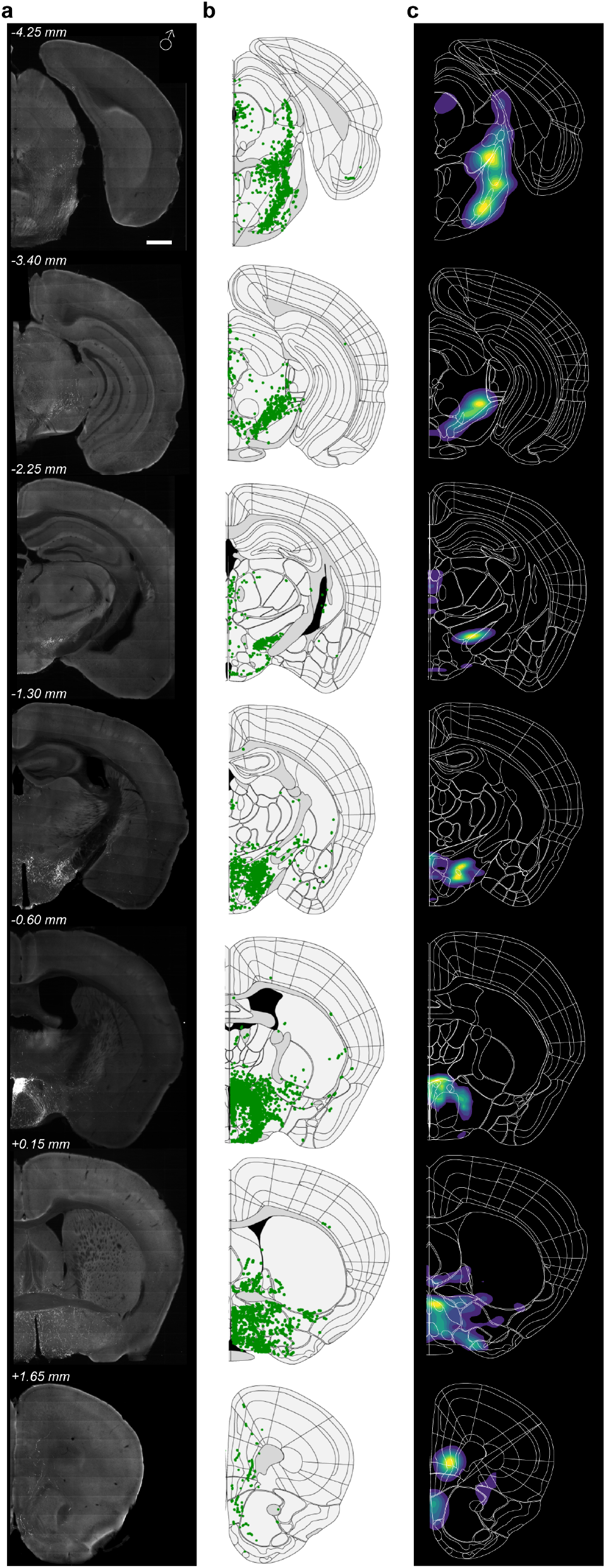
Oxt+ axonal projections are broadly distributed in males. a. Raw images of axonal projections of Oxt+ paraventricular neurons from a representative male mouse. Scale bar, 0.5 mm. b. Detected axon pixels using Sobel edge detection algorithm of images shown in panel a. Brain slices were registered to Allen brain atlas slice with a modified version of the WholeBrain software package. c. Density contour of axonal projections showing nuclei with highest normalized pixel count.

**Figure 1—Figure Supplement 3:**
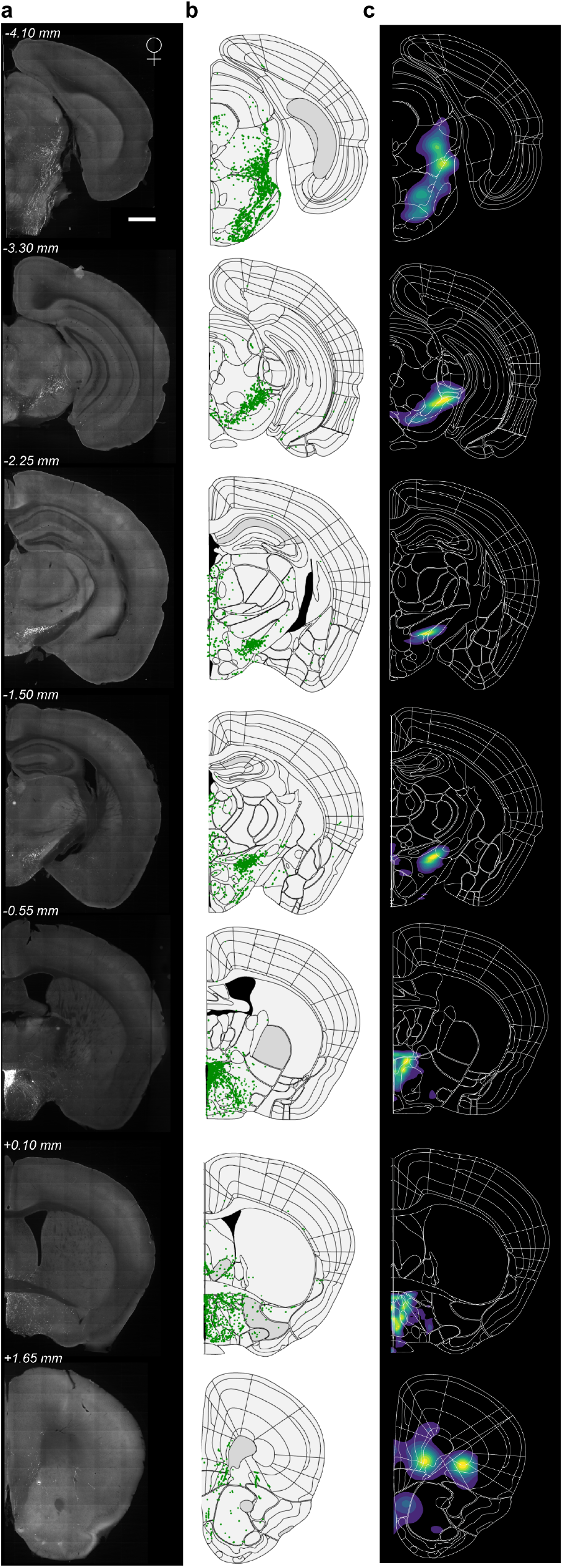
Oxt+ axonal projections are broadly distributed in females. a. Raw representative images of Oxt+ axons in a female mouse. Scale bar, 0.5 mm. b. Segmented Oxt+ fibers using Sobel edge detection algorithm and registered to Allen reference atlas. c. Density contour plots showing brain regions of high pixel density.

**Figure 1—Figure Supplement 4:**
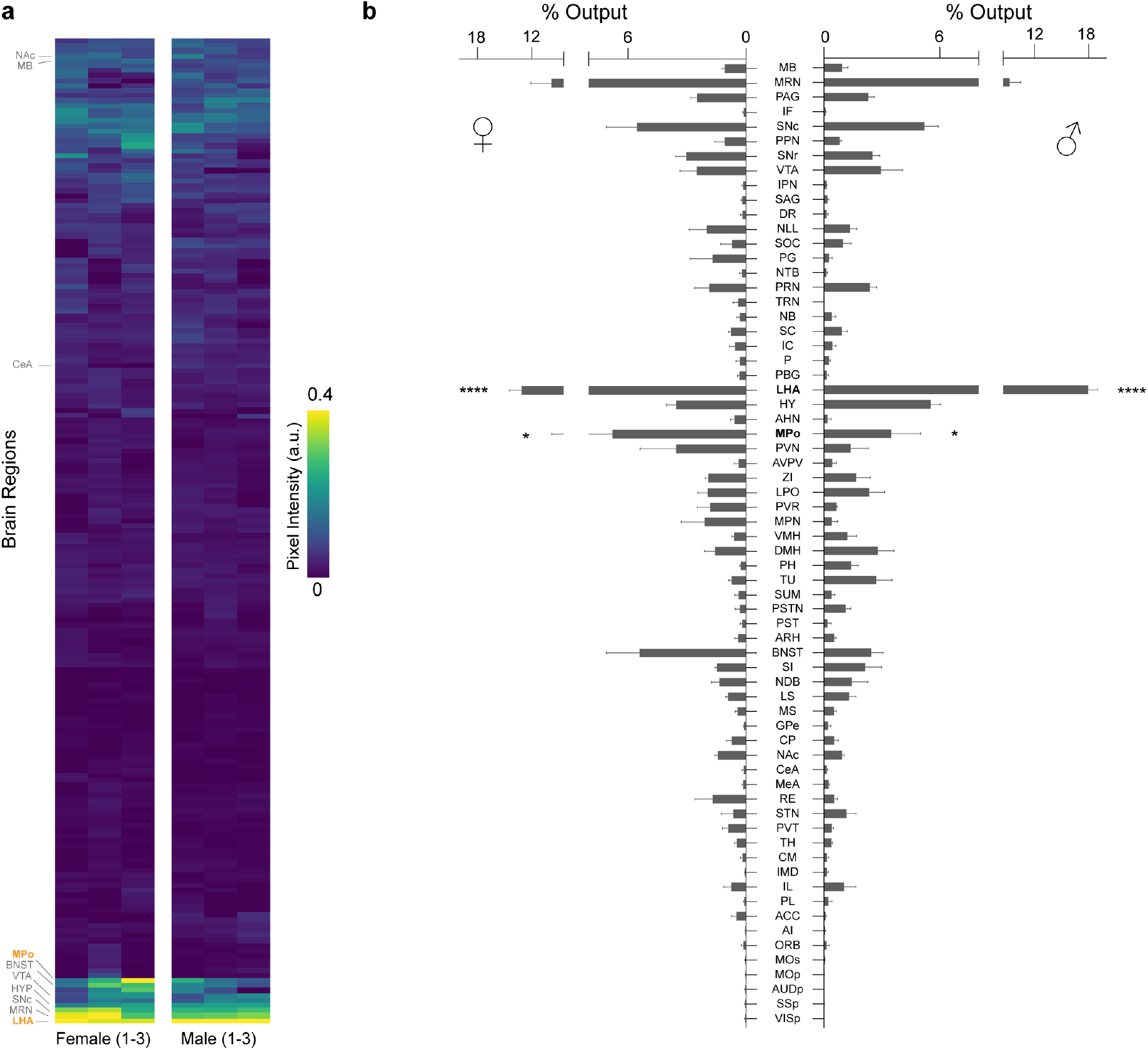
Oxt+ axonal projections are largely sex-invariant. a. Heatmap of all brain regions identified as containing Oxt+ axonal fibers. Each column represents one sample and each row represents a given brain region. Pixel counts were normalized for all samples and depicted as a heatmap with yellow marking areas with prominent Oxt+ fiber labeling and dark purple regions representing areas with little to no Oxt+ axons. Overall, males and females exhibit similar axonal patterns for nearly all nuclei in Oxt dataset. Regions labeled in bold orange font are statistically significant. b. Bar graph comparing regions with Oxt fibers in males (n=3) and females (n=3). Bar graph includes regions with Oxt labeling of >0.15% of total output. For a complete list of regions and quantification, see Table 1. LHA: **** adjusted p< 0.0001, MPo: * adjusted p=0.0113, two-way ANOVA using Bonferroni post hocs for multiple comparisons.

**Figure 2—Figure Supplement 1:**
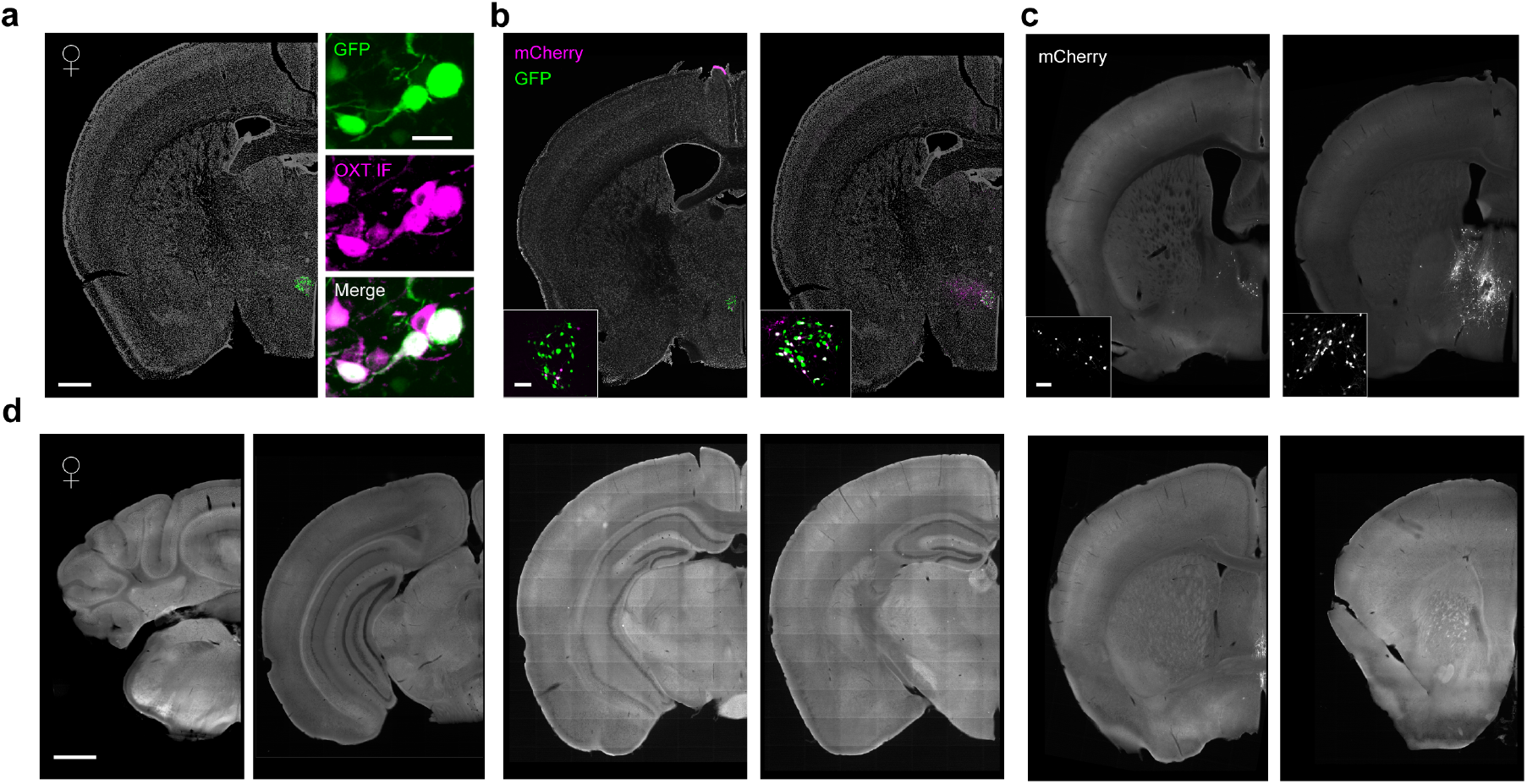
PVN Oxt+ inputs are not observed in distal midbrain or forebrain nuclei. a. *Left* Representative coronal slice of Oxt-cre female injected with Cre-dependent AAV-H2B-GFP-2A-oG, AAV-EF1a-GT (helper virus). GFP expression (green) is observed in the PVN, overlayed with DAPI stain (gray). Scale bar, 0.5 mm. *Right* GFP expression is restricted to PVN cells immunostained with anti-Oxt antibody. Scale bar, 40 µm. b. *Left* Image of female Oxt-cre mouse that had few (<5 cells) cells co-transfected with rabies and helper virus. This sample was excluded from quantitative group analyses and served as a negative control compared to samples with co-transfection of both rabies and helper virus to allow for retrograde tracing, depicted in the *right* panel. mCherry cells transfected with rabies virus are depicted in magenta and cells with GFP expression are depicted in green. Scale bar, 40 µm. c. Matched example slices with mCherry+ cells retrogradely labeled from the samples depicted in panel b. Sample on the left with minimal starter cells shows limited expression of mCherry and was excluded from analysis. By comparison, the sample on the right shows more prominent retrograde labeling. Scale bar, 40 µm. d. Example coronal images from sample depicted in panel a. Oxt neurons show no labeling of distal inputs from cerebellar, hindbrain, midbrain, or cortical regions. Scale bar, 0.5 mm.

**Figure 3—Figure Supplement 1:**
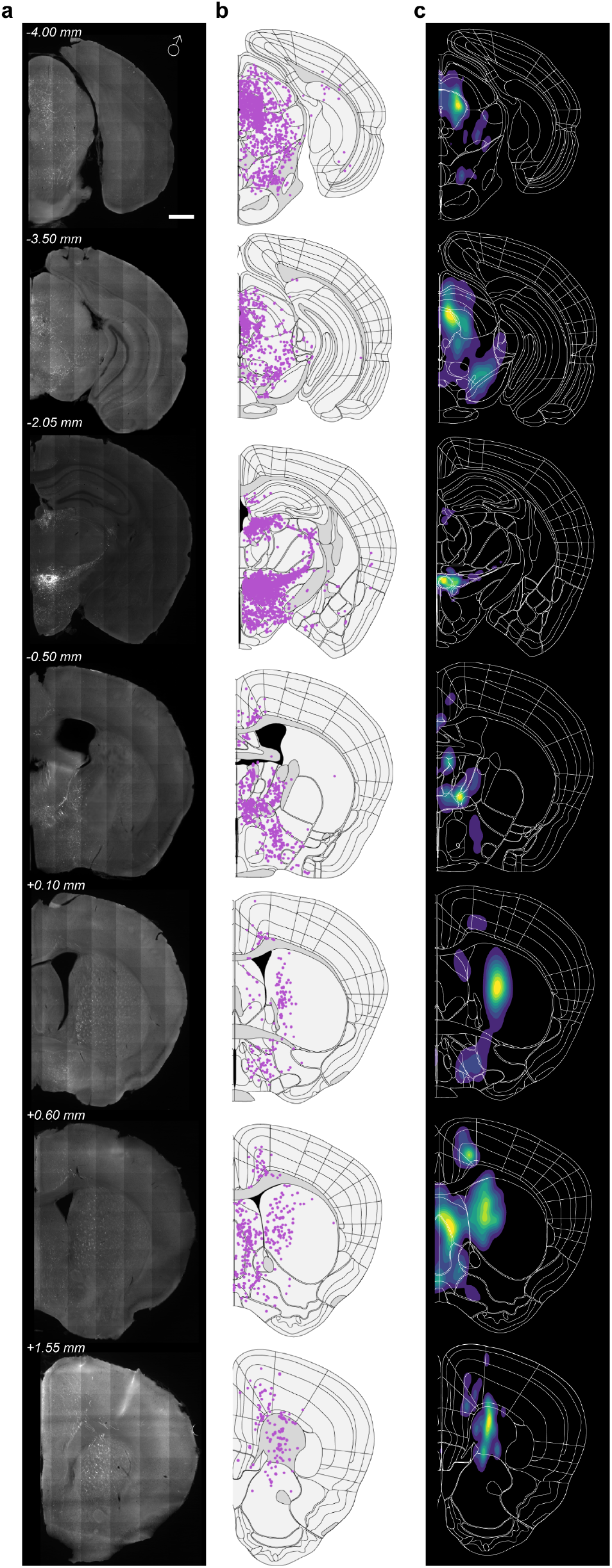
Avp+ projections in male mice densely label hypothalamic and midbrain nuclei. a. Raw representative images of Avp+ axons in a male mouse. Male mice show eYFP+ cell labeling in extra-paraventricular nuclei, including BNST and ZI that surround PVN, Scale bar 0.5 mm. b. Detected and registered Avp+ fibers using Sobel edge detection algorithm. c. Density contour of axonal projections showing nuclei with highest normalized pixel counts.

**Figure 3—Figure Supplement 2:**
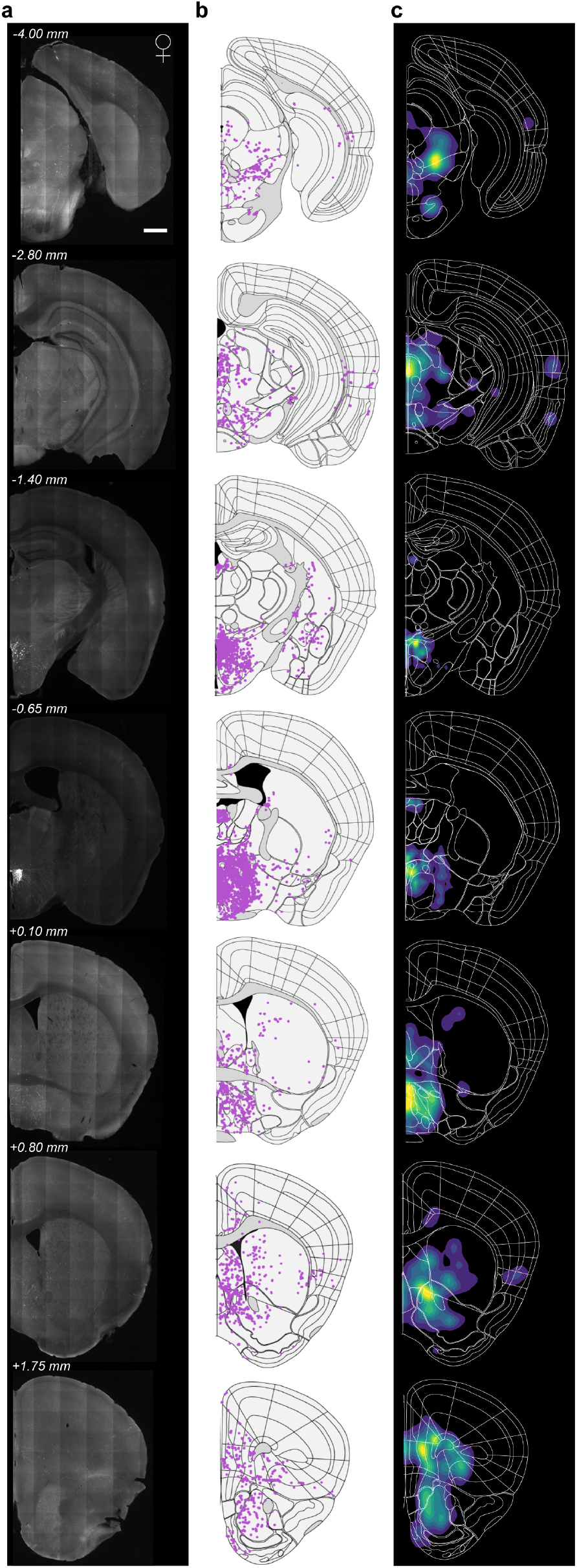
Avp+ PVN neuron projections in females extend to distal cortical and striatal nuclei. a. Raw representative images of Avp+ axons in a female mouse. Somatic eYFP+ labeling more restricted to PVN, compared to male Avp-cre mice. Scale bar, 0.5 mm. b. Detected and registered Avp+ axons using Sobel edge detection algorithm. c. Density contour of axonal projections showing nuclei with highest normalized pixel counts within regions of interest.

**Figure 3—Figure Supplement 3:**
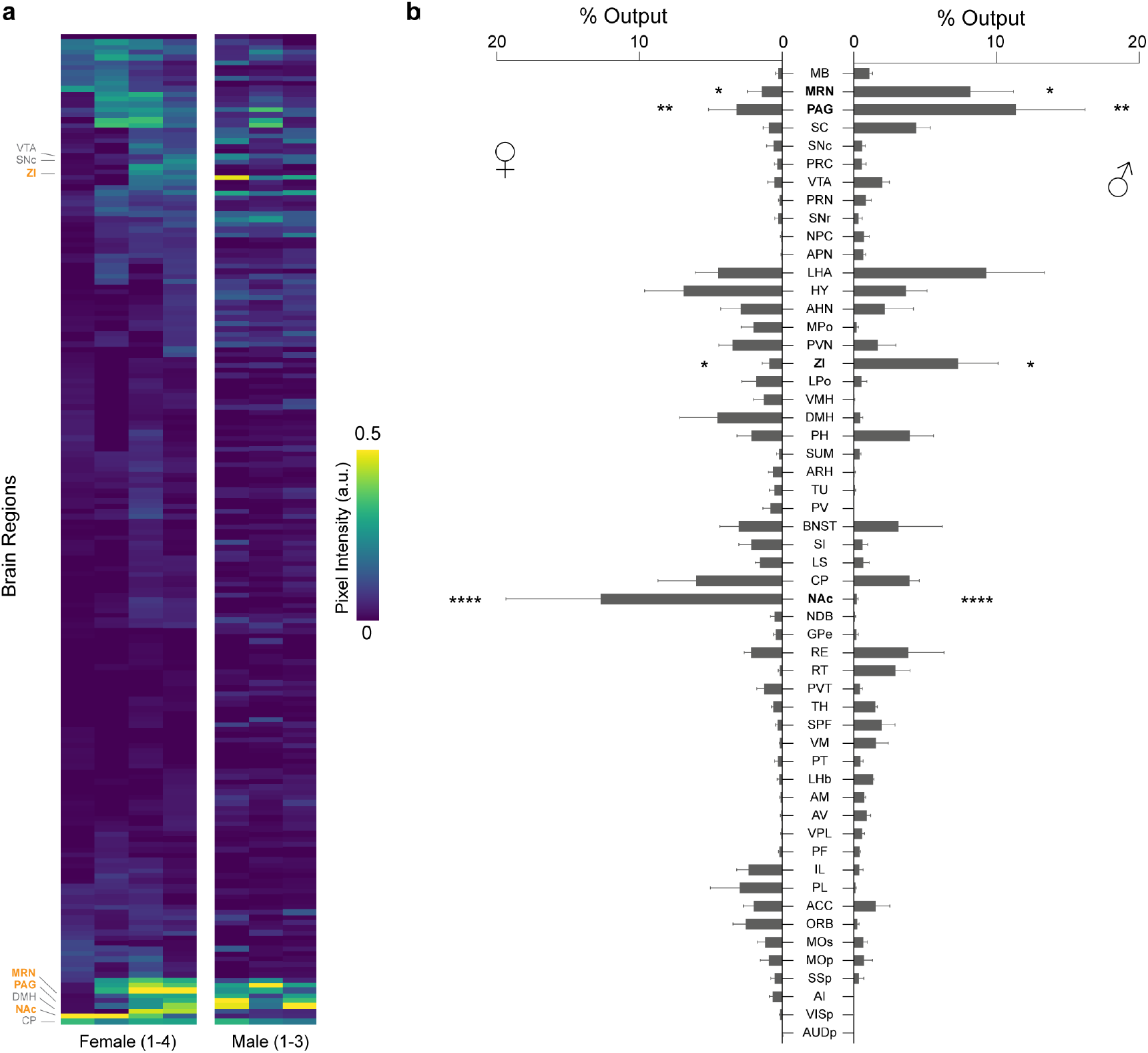
Avp+ PVN projections exhibit sex-dependent distributions. a. Heatmap of all brain regions identified to contain Avp+ axonal fibers, each column represents one sample and each row represents a given brain region. Pixel counts were normalized for all samples and depicted as a heatmap with yellow depicting areas with robust Avp+ fiber labeling and dark purple regions representing areas with little to no Avp+ axons. Regions labeled with bold orange font are statistically significant. b. Bar graph comparing regions with Avp projections across females (n=4) and males (n=3). Regions displayed in the bar graph exhibited Avp labeling >0.2% of total output. For a complete list of regions and quantification, see Table 2. MRN: * adjusted p-value= 0.0235, PAG: ** adjusted p-value = 0.0011, ZI: adjusted p-value=0.0405, NAc: **** adjusted p-value <0.0001, two-way ANOVA using Bonferroni post hocs for multiple comparisons.

**Figure 4—Figure Supplement 1:**
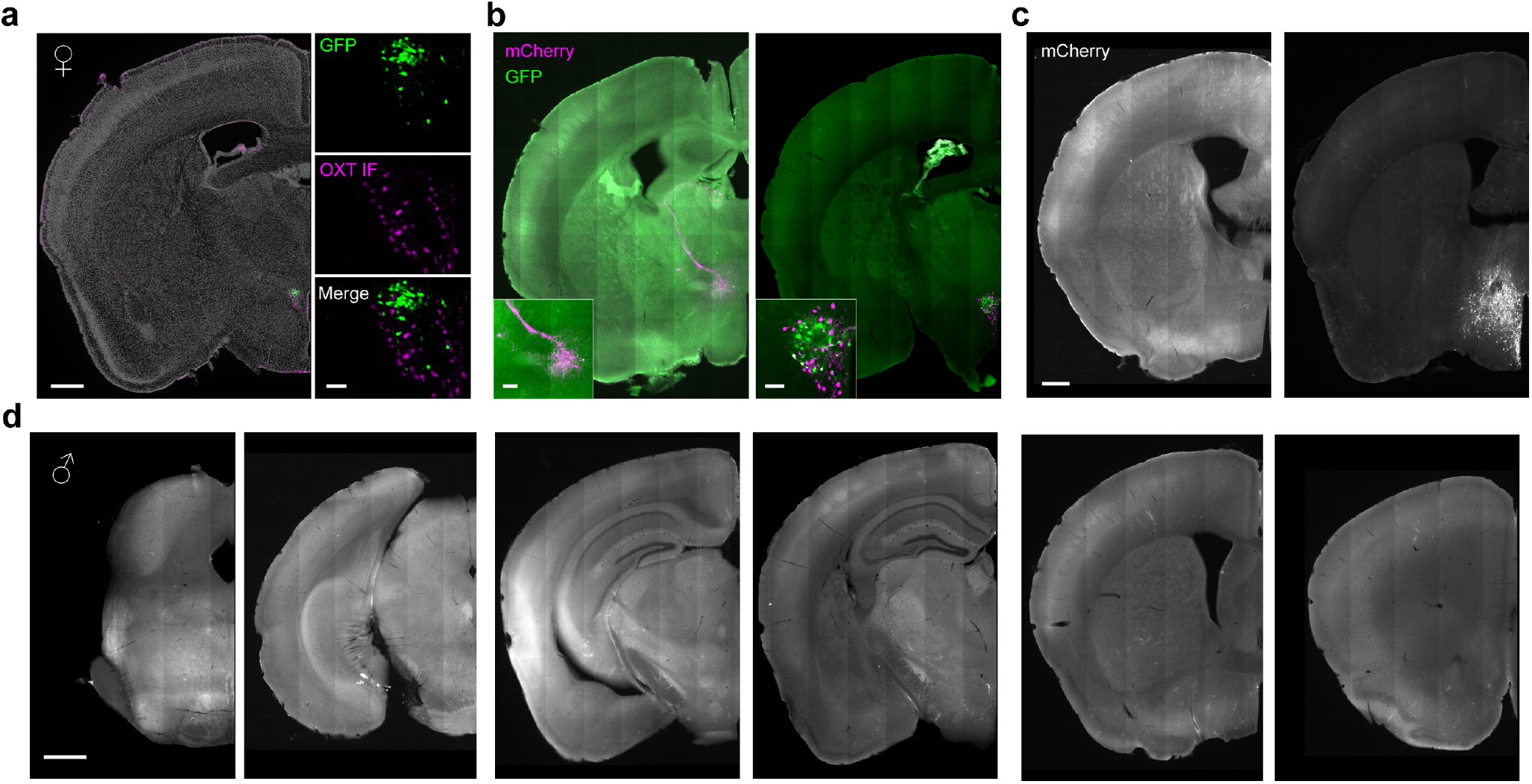
PVN Avp+ inputs are not observed in distal midbrain or forebrain nuclei. *a. Left* Representative coronal slice of Avp-cre female injected with Cre-dependent AAV-H2B-GFP-2A-oG, AAV-EF1a-GT (helper virus). GFP expression (green) is observed in the PVN, overlayed with DAPI stain (gray) and shows little overlap with cells immunostained with anti-Oxt antibody (magenta). Scale bar, 0.5mm. *Right* GFP expression is restricted to PVN Avp+ cells, but not observed in Oxt+ PVN cells. Scale bar, 100 µm. *b. Left* Image of female Avp-cre mouse that had no GFP + cells in the PVN and minimal ectopic expression of rabies (mCherry) virus. Ectopic expression of rabies mCherry signal was qualitatively different than samples co-transfected with helper and rabies virus. This sample was excluded from quantitative group analyses and served as a negative control compared to samples with sufficient starter cell populations, depicted in the *right* panel. mCherry cells transfected with rabies virus are depicted in magenta and cells with GFP expression are depicted in green. Scale bar, 40 µm. *c.* Matched example slices with mCherry+ cells retrogradely labeled from the samples depicted in panel b. Sample on the left with no starter cells shows no mCherry signal in extra-hypothalamic nuclei and was excluded from analysis. By comparison, the sample on the right shows more robust retrograde labeling. Scale bar, 0.5 mm. *d.* Example coronal images extending from hindbrain to forebrain of sample depicted in Figure 4. Avp PVN neurons show no labeling of distal inputs from cerebellar, hindbrain, midbrain, or cortical regions. Scale bar, 0.5 mm.

## References

1. Baribeau, D. A. & Anagnostou, E. Oxytocin and vasopressin: Linking pituitary neuropeptides and their receptors to social neurocircuits. Front. Neurosci. 9, 335 (2015).

2. Carter, C. S. The Oxytocin–Vasopressin Pathway in the Context of Love and Fear. Front. Endocrinol. 8, 356 (2017).

3. Huber, D., Veinante, P. & Stoop, R. Vasopressin and oxytocin excite distinct neuronal populations in the central amygdala. Science 308, 245–248 (2005).

4. Donaldson, Z. R. & Young, L. J. Oxytocin, vasopressin, and the neurogenetics of sociality. Science 322, 900–904 (2008).

5. Rickenbacher, E., Perry, R. E., Sullivan, R. M. & Moita, M. A. Freezing suppression by oxytocin in central amygdala allows alternate defensive behaviours and mother-pup interactions. Elife 6, e24080 (2017).

6. Marlin, B. J., Mitre, M., D’amour, J. A., Chao, M. V. & Froemke, R. C. Oxytocin enables maternal behaviour by balancing cortical inhibition. Nature 520, 499–504 (2015).

7. Nishimori, K. et al. Oxytocin is required for nursing but is not essential for parturition or reproductive behavior. Proc. Natl. Acad. Sci. U. S. A. 93, 11699–11704 (1996).

8. Nakajima, M., Görlich, A. & Heintz, N. Oxytocin modulates female sociosexual behavior through a specific class of prefrontal cortical interneurons. Cell 159, 295–305 (2014).

9. Li, K., Nakajima, M., Ibañez-Tallon, I. & Heintz, N. A Cortical Circuit for Sexually Dimorphic Oxytocin-Dependent Anxiety Behaviors. Cell 167, 60-72.e11 (2016).

10. Melis, M. R., Succu, S., Sanna, F., Boi, A. & Argiolas, A. Oxytocin injected into the ventral subiculum or the posteromedial cortical nucleus of the amygdala induces penile erection and increases extracellular dopamine levels in the nucleus accumbens of male rats. Eur. J. Neurosci. 30, 1349–1357 (2009).

11. Calcagnoli, F. et al. Oxytocin microinjected into the central amygdaloid nuclei exerts anti-aggressive effects in male rats. Neuropharmacology 90, 74–81 (2015).

12. Oti, T. et al. Oxytocin Influences Male Sexual Activity via Non-synaptic Axonal Release in the Spinal Cord. Curr. Biol. 31, 103-114.e5 (2021).

13. Winslow, J. T., Hastings, N., Carter, C. S., Harbaugh, C. R. & Insel, T. R. A role for central vasopressin in pair bonding in monogamous prairie voles. Nature 365, 545–548 (1993).

14. Nair, H. P. & Young, L. J. Vasopressin and pair-bond formation: Genes to brain to behavior. Physiology 21, 146–152 (2006).

15. Carter, C. S. & Perkeybile, A. M. The Monogamy Paradox: What do love and sex have to do with it? Front. Ecol. Evol. 6, 202 (2018).

16. Knobloch, H. S. et al. Evoked axonal oxytocin release in the central amygdala attenuates fear response. Neuron 73, 553–566 (2012).

17. Grinevich, V. & Stoop, R. Interplay between Oxytocin and Sensory Systems in the Orchestration of Socio-Emotional Behaviors. Neuron 99, 887–904 (2018).

18. Tang, Y. et al. Social touch promotes interfemale communication via activation of parvocellular oxytocin neurons. Nat. Neurosci. 23, 1125–1137 (2020).

19. Carcea, I. et al. Oxytocin neurons enable social transmission of maternal behaviour. Nature 596, 553–557 (2021).

20. Mitre, M. et al. A Distributed Network for Social Cognition Enriched for Oxytocin Receptors. J. Neurosci. 36, 2517–2535 (2016).

21. Dumais, K. M. & Veenema, A. H. Vasopressin and oxytocin receptor systems in the brain: Sex differences and sex-specific regulation of social behavior. Front. Neuroendocrinol. 40, 1–23 (2016).

22. Fürth, D. et al. An interactive framework for whole-brain maps at cellular resolution. Nat. Neurosci. 21, 139–149 (2018).

23. Lein, E. S. et al. Genome-wide atlas of gene expression in the adult mouse brain. Nature 445, 168–176 (2007).

24. Shah, B. P. et al. MC4R-expressing glutamatergic neurons in the paraventricular hypothalamus regulate feeding and are synaptically connected to the parabrachial nucleus. Proc. Natl. Acad. Sci. 111, 13193–13198 (2014).

25. Wu, Z. et al. An Obligate Role of Oxytocin Neurons in Diet Induced Energy Expenditure. PLoS One 7, e45167 (2012).

26. Xiao, L., Priest, M. F. & Kozorovitskiy, Y. Oxytocin functions as a spatiotemporal filter for excitatory synaptic inputs to VTA dopamine neurons. Elife 7, e33892 (2018).

27. Li, C. et al. Defined Paraventricular Hypothalamic Populations Exhibit Differential Responses to Food Contingent on Caloric State. Cell Metab. 29, 681-694.e5 (2019).

28. Sun, W. et al. Oxytocin Relieves Neuropathic Pain Through GABA Release and Presynaptic TRPV1 Inhibition in Spinal Cord. Front. Mol. Neurosci. 11, 248 (2018).

29. Marlin, B. J., Mitre, M., D’amour, J. A., Chao, M. V. & Froemke, R. C. Oxytocin enables maternal behaviour by balancing cortical inhibition. Nature 520, 499–504 (2015).

30. Schiavo, J. K. et al. Innate and plastic mechanisms for maternal behaviour in auditory cortex. Nature 587, 426–431 (2020).

31. Wei, Y.-C. et al. Medial preoptic area in mice is capable of mediating sexually dimorphic behaviors regardless of gender. Nat. Commun. 9, 279 (2018).

32. Tasker, J. G., Boudaba, C. & Schrader, L. A. Local Glutamatergic and GABAergic Synaptic Circuits and Metabotropic Glutamate Receptors in the Hypothalamic Paraventricular and Supraoptic Nuclei. Adv. Exp. Med. Biol. 449, 117–121 (1998).

33. Boudaba, C., Szabó, K. & Tasker, J. G. Physiological Mapping of Local Inhibitory Inputs to the Hypothalamic Paraventricular Nucleus. J. Neurosci. 16, 7151 (1996).

34. Roland, B. L. & Sawchenko, P. E. Local origins of some GABAergic projections to the paraventricular and supraoptic nuclei of the hypothalamus in the rat. J. Comp. Neurol. 332, 123–143 (1993).

35. Silverman, A. J., Hoffman, D. L. & Zimmerman, E. A. The descending afferent connections of the paraventricular nucleus of the hypothalamus (PVN). Brain Res. Bull. 6, 47–61 (1981).

36. Cservenák, M. et al. A Thalamo-Hypothalamic Pathway That Activates Oxytocin Neurons in Social Contexts in Female Rats. Endocrinology 158, 335–348 (2017).

37. Fakhoury, M., Salman, I., Najjar, W., Merhej, G. & Lawand, N. The Lateral Hypothalamus: An Uncharted Territory for Processing Peripheral Neurogenic Inflammation. Front. Neurosci. 14, 101 (2020).

38. Bonnavion, P., Mickelsen, L. E., Fujita, A., de Lecea, L. & Jackson, A. C. Hubs and spokes of the lateral hypothalamus: cell types, circuits and behaviour. J. Physiol. 594, 6443–6462 (2016).

39. Xiao, L., Priest, M. F., Nasenbeny, J., Lu, T. & Kozorovitskiy, Y. Biased Oxytocinergic Modulation of Midbrain Dopamine Systems. Neuron 95, 368-384.e5 (2017).

40. Harris, J. A., Hirokawa, K. E., Sorensen, S. A., Gu, H. & Mills, M. Anatomical characterization of Cre driver mice for neural circuit mapping and manipulation. Front Neural Circuits 8, 76 (2014).

41. Duque-Wilckens, N. et al. Extrahypothalamic oxytocin neurons drive stress-induced social vigilance and avoidance. Proc. Natl. Acad. Sci. U. S. A. 117, 26406–26413 (2020).

42. Kelly, A. M., Hiura, L. C., Saunders, A. G. & Ophir, A. G. Oxytocin Neurons Exhibit Extensive Functional Plasticity Due To Offspring Age in Mothers and Fathers. Integr. Comp. Biol. 57, 603–618 (2017).

43. Rigney, N., Whylings, J., Mieda, M., De Vries, G. J. & Petrulis, A. Sexually Dimorphic Vasopressin Cells Modulate Social Investigation and Communication in Sex-Specific Ways. eNeuro 6, 0415–0418 (2019).

44. DiBenedictis, B. T., Nussbaum, E. R., Cheung, H. K. & Veenema, A. H. Quantitative mapping reveals age and sex differences in vasopressin, but not oxytocin, immunoreactivity in the rat social behavior neural network. J. Comp. Neurol. 525, 2549–2570 (2017).

45. Silverman, A.-J. & Oldfield, B. J. Synaptic input to vasopressin neurons of the paraventricular nucleus (PVN). Peptides 5, 139–150 (1984).

46. Wei, H. H. et al. Presynaptic inputs to vasopressin neurons in the hypothalamic supraoptic nucleus and paraventricular nucleus in mice. Exp. Neurol. 343, 113784 (2021).

47. Dumais, K. M. & Veenema, A. H. Vasopressin and oxytocin receptor systems in the brain: Sex differences and sex-specific regulation of social behavior. Front. Neuroendocrinol. 40, 1–23 (2016).

48. Herman, J. P., Tasker, J. G., Ziegler, D. R. & Cullinan, W. E. Local circuit regulation of paraventricular nucleus stress integration: Glutamate–GABA connections. Pharmacol. Biochem. Behav. 71, 457–468 (2002).

49. Fakhoury, M., Salman, I., Najjar, W., Merhej, G. & Lawand, N. The Lateral Hypothalamus: An Uncharted Territory for Processing Peripheral Neurogenic Inflammation. Front. Neurosci. 14, 101 (2020).

50. Roland, B. L. & Sawchenko, P. E. Local origins of some GABAergic projections to the paraventricular and supraoptic nuclei of the hypothalamus in the rat. J. Comp. Neurol. 332, 123–143 (1993).

51. Grinevich, V. & Stoop, R. Review Interplay between Oxytocin and Sensory Systems in the Orchestration of Socio-Emotional Behaviors. Neuron 99, 887–904 (2018).

52. Mitre, M., Minder, J., Morina, E. X., Chao, M. V. & Froemke, R. C. Oxytocin Modulation of Neural Circuits. in Current Topics in Behavioral Neurosciences 35, 31–53 (2018).

